# Functional relevance of mobile and clustered Ca_V_2.1 channels in central synapses

**DOI:** 10.64898/2026.07.13.737722

**Authors:** El Khallouqi Abderazzaq, Carolina Amaral, Anna Schroder Lassen, Stephan Weißbach, Corinna Werkmann, Arthur Bikbaev, Melanie D. Mark, Stefan Herlitze, Jennifer Heck, Alexander M. Walter, Martin Heine

**Affiliations:** Functional Neurobiology, Institute of Developmental Biology and Neurobiology, Johannes Gutenberg University Mainz, Mainz 55128, Germany; Behavioural Neuroscience, Ruhr-University Bochum, Bochum 44780, Germany; Molecular Neurophysiology, Department of Neuroscience, Faculty of Health and Medical Sciences, University of Copenhagen, Copenhagen 2200, Denmark; Genome Biology Unit, European Molecular Biology Laboratory, Heidelberg, Germany; Department of Zoology and Neurobiology, Ruhr-University Bochum, Bochum, Germany

## Abstract

Reliable neurotransmitter release critically depends on the spatial relationship between voltage-gated calcium channels (VGCCs) and presynaptic release sites. Single particle tracking of endogenous Ca_V_2.1 channels at glutamatergic synapses of hippocampal neurons revealed that apart from Ca_V_2.1 channels aggregated in stable nanodomain clusters, a substantial fraction of Ca_v_2.1 channels remained mobile, raising the question of whether these dispersed channels contribute to synaptic function. Mathematical modelling predicted that dispersed Ca_v_2.1 channels cooperatively enhance release reliability. Upon repetitive stimulation, mobile Ca_V_2.1 channels enable alternative use of release sites and thereby reduce the probability of failed presynaptic release. Both optogenetic immobilisation of Ca_V_2.1 channels *per se* or activation of GABA_B_ receptors (GABA_B_Rs) alone increase the failure rate and can lead to synaptic silencing. However, optogenetic clustering Ca_V_2.1 channels prior to GABA_B_R activation increases the fraction of synapses that remain active even in presence of GABA_B_R agonist. The contribution of mobile channels to reliable neurotransmitter release is frequency-dependent and is minor at stimulation frequencies 1 Hz but becomes strong at frequencies over 10 Hz. These results demonstrate that mobile presynaptic Ca_V_2.1 channels increase the frequency range of synaptic transmission but are particularly sensitive to metabotropic GABA_B_R-mediated inhibition in glutamatergic hippocampal synapses.

## Introduction

The importance of Ca_V_2.1 channels in triggering presynaptic neurotransmitter release has been extensively demonstrated using functional and structural approaches. Rapid but very transient opening of Ca_V_2.1 channels upon presynaptic depolarization, in conjunction with low affinity vesicular Ca^2+^ sensors or abundance of intracellular Ca^2+^ buffers, leads to Ca^2+^ influx that is strongly restricted both in time and space. Therefore, a reliable neurotransmitter release demands positioning of channels in close proximity to synaptic vesicles (SV) at the release site. An additional approach to increase the release probability (P_r_) is clustering of Ca_V_2.1 channels that can partially counteract spatiotemporal limitations of Ca^2+^ influx. Rather clustered than a random distribution of calcium channels tightly linked to ready releasable vesicles was demonstrated in several synapses under control conditions and was found to be disturbed under pathophysiological conditions (Holderith et al., 2012; Kim et al., 2024; Martín-Belmonte et al., 2025; Miki et al., 2017; Nakamura et al., 2015; Rebola et al., 2019). Several identified direct and indirect interaction sites of the intracellular β-subunit and the pore-forming α_1_ subunit of Ca_V_2.1 and Ca_V_2.2 channels with presynaptic scaffold proteins contribute to formation of synaptic channel clusters and potentially involve mechanisms based on liquid-liquid phase separation (Davydova et al., 2014; Emperador-Melero et al., 2024; Kaeser et al., 2011; Wu et al., 2019). However, studies with deletion or point mutations of protein-protein interactions in the C terminus of the α_1_ subunit revealed synaptic accumulation of VGCCs despite the loss from a substantial fraction of interaction sites (Chin and Kaeser, 2024; Heck et al., 2019; Hu et al., 2005; Li et al., 2024; Lubbert et al., 2017). These data suggests that calcium channels lacking tight associations with releasable vesicles can still be targeted to active zones and contribute to synaptic release.

But how flexible are the links between release sites and calcium channels within the presynaptic membrane? Does this have direct consequences for the release probability of a given synapse? Most glutamatergic synapses of cortical and hippocampal synapses display an extremely wide range of release probabilities (for review see (Branco and Staras, 2009)) and have only one active zone with several release sites (Maschi and Klyachko, 2017; Maschi and Klyachko, 2020). Several parameters, including the number of VGCCs and the size of ready releasable pool, can underlie such diversity in the Pr. Importantly, a fraction of presynaptic calcium channels is mobile, which is not surprising given the large rearrangements of the presynaptic membrane during vesicle fusion and retrieval (Byczkowicz et al., 2018). We and others have previously shown that Ca_V_2 channels are indeed not statically distributed in the active zone but are able to jump over > 50 nm distances within tens of milliseconds (Heck *et al*., 2019; Mercer et al., 2011; Schneider et al., 2015). Given relative stability of release sites, such mobility of VGCCs would cause an ongoing fluctuation between their tight or loose coupling to SVs that can dynamically affect the P_r_ of a given synapse. Furthermore, the surprisingly fast spatial dynamics of calcium channels within the presynaptic membrane raises the question whether these channels participate in SV release or should be considered as non-functional.

Using a knock-in mouse model of N-terminal citrine-tagged Ca_V_2.1 channels (Mark et al., 2011), we developed an approach to localize and monitor endogenous Ca_V_2.1 channels over time, as well as to correlate their presynaptic position with recently used release sites. In line with earlier data, we found that clustered channels generally represent only a fraction in hippocampal synapses, and the number of clustered channels is highly variable between presynaptic boutons even along the same axon. Interference with the local dynamics of Ca_V_2.1 channels by optogenetic clustering of the channels resulted in a less robust transmitter release particular at frequencies >10 Hz and reduced the engagement of release sites within the presynaptic membrane. Modulation of the presynaptic release probability is robustly controlled by various metabotropic receptors within the presynaptic membrane. We found that pharmacological activation of GABA_B_Rs led to silencing of most glutamatergic synapses. However, one third of presynaptic terminals, where clustered Ca_V_2.1 channels were prevalent, remained active and maintained evoked glutamate release even in presence of GABA_B_R agonist. Pre-clustering of Ca_V_2.1 channels prevented GABA_B_R-mediated silencing of synapses, indicating that the mobile Ca_V_2.1 channels are the primary target of G protein-mediated neuromodulation. Hence, the robustness of evoked neurotransmitter release depends on a clustered fraction of Ca_V_2.1 channels, whereas alternative engagement of all available release sites upon high-frequency activation is defined by the mobile population of presynaptic Ca_V_2.1 channels.

## Results

### Tagging of endogenous Ca_V_2.1 channels reveals their ffexible synaptic residence

Changes in the number, distribution, kinetic properties or expression of different Ca_V_2 isoforms in the synaptic membrane have significant impact on the release probability and replenishment (Thanawala and Regehr, 2013; Wang and Kaczmarek, 1998)) of synaptic vesicles, as well on homeostatic plasticity of synapses (Cao and Tsien, 2010; Chin and Kaeser, 2024; Fedchyshyn and Wang, 2005; Hefft and Jonas, 2005; Held et al., 2020; Krick et al., 2021; Lübbert et al., 2019; Reid et al., 2003). To directly correlate organization of endogenous presynaptic Ca_V_2.1 channels with synaptic function, first we performed their fluorescent labelling. N-terminally tagged endogenous Ca_V_2 used previously in different models (Gratz et al., 2019; Mark *et al*., 2011; Mueller et al., 2023) did not affect their kinetic properties or synaptic localization (Gratz *et al*., 2019; Mark *et al*., 2011). Other tagging approaches using mEOS or HaloTag were also reported to be functionally neutral (Ghelani et al., 2023; Medeiros et al., 2024; Schneider *et al*., 2015). Here, we used the mouse model introduced by Mark et al. (2011) with an insertion of citrine at the N-terminus of Ca_V_2.1 channels. Despite the monomeric nature of the citrine tag, the autofluorescence of neurons and glial cells limited usability of the tag as a reliable reporter of spatial location of individual Ca_V_2.1 channels. To overcome this limitation, we used the citrine tag as epitope for a linker protein that is based on anti-GFP nanobodies. To verify the specificity of anti-GFP nanobodies, we compared signals of Ca_V_2.1 channels labelled using N-terminal nanobody and C-terminal antibody and found nearly complete overlap (Suppl.Fig. 1A-C). Cre recombinase-dependent knock-down (KD) of Ca_V_2.1 channels confirmed the specificity of both labelling approaches (Suppl.Fig. 1A-C). Fusion of HaloTag to the intrabody against GFP allowed us to use cell permeable fluorescent HaloTag ligands to label individual Ca_V_2.1 channels and follow their distribution over time. The covalent binding of the HaloTag ligand preserves the 1:1 stoichiometry between fluorescent label and channel (Encell et al., 2012; Wilhelm et al., 2021). Hence, fusion of HaloTag to the intrabody against GFP enabled the use of cell-permeable fluorescent HaloTag ligands to label individual Ca_V_2.1 channels and monitor their location over time. The cytosolic expression of the intrabody was controlled by a transcription feedback loop (CCR5/KRAB(A)) (Gross et al., 2013) to restrict the number of free cytosolic unbound intrabodies. Several linker sequences between intrabody and the HaloTag have been tested to optimise the labelling efficiency. For testing the targeting efficiency of the intrabody, we used HEK293T cells expressing the intrabody together with a GFP targeted to the outer (GPI-GFP) or inner (MyrGFP) leaflet of the cellular membrane were used to access the targeting efficiency of the intrabody. We observed a clear nuclear localization of the intrabody if GFP was extracellularly expressed. Co-expression of a myristoylated GFP confirmed a defined targeting of the intrabody to the inner plasma membrane (Suppl. Fig. 1D, E). In cultured hippocampal neurons, we found a targeted expression of the intrabody under control conditions, while strong nuclear expression of the intrabody was evident upon Cre recombinase-mediated KD of Ca_V_2.1 channels at DIV 3 *in vitro* (Fig. 1D, I). For the intrabody lacking CCR5/KRAB(A), we observed a massive cytosolic expression of the intrabody (Fig. 1F, I).

**Figure 1:**
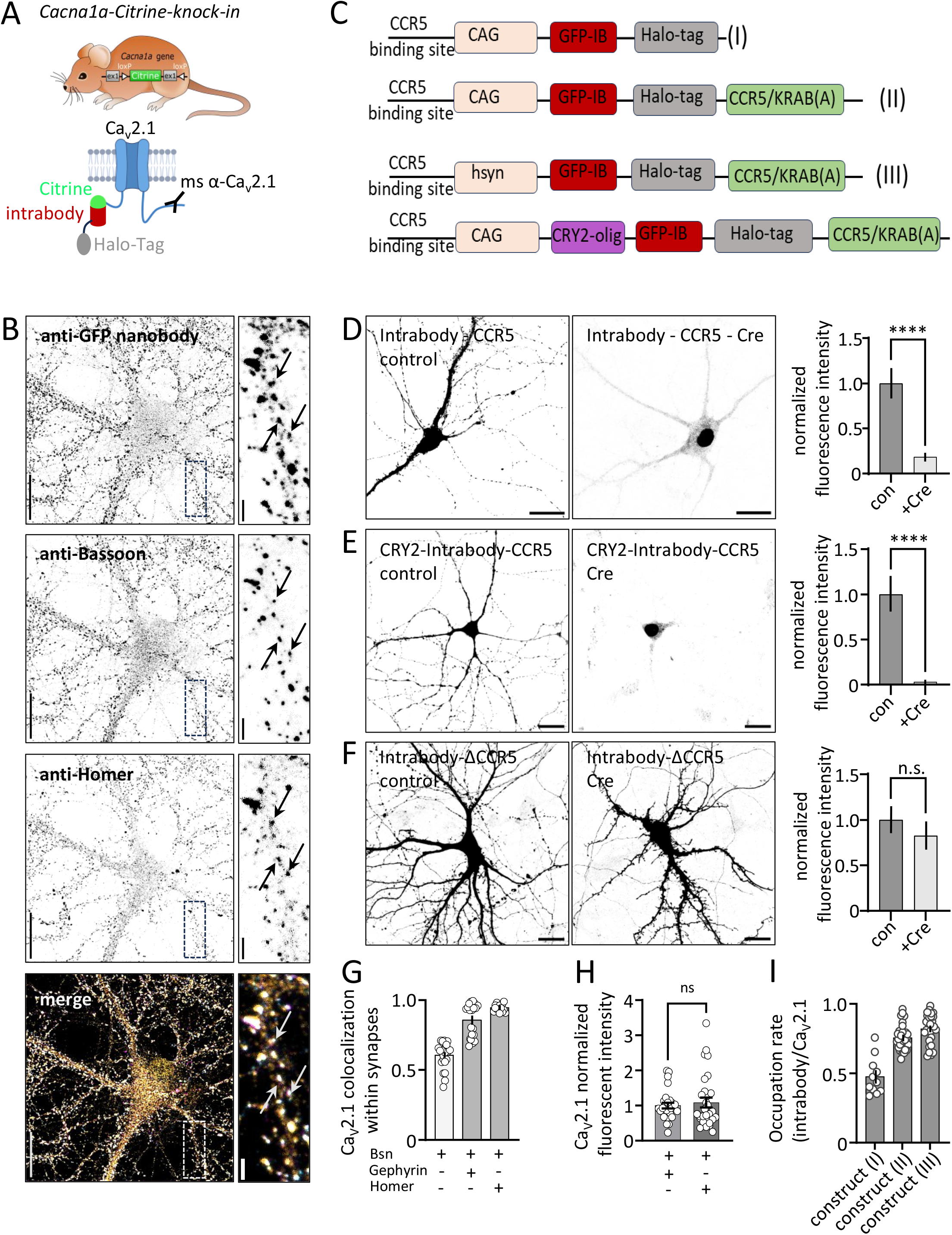
Labelling of the endogenous population of Ca_V_2.1 channels in hippocampal neurons, using the expression of an intrabody under transcription control. **A)** Used mouse model for labelling the endogenous population of Cav2.1 channels and the strategy how to use the N-terminal citrine-tag as defined epitope. The binding site of antibodies on the C-terminus is indicated as well, which allowed us to control the specificity of intrabody labelling. **B)** Hippocampal neurons 16 DIV from the Cacna1a^Citrine^ KI mouse labelled with anti-GFP nanobody (ATTO647N), anti-bassoon as presynaptic scaffold, and anti-Homer as postsynaptic scaffold protein. Within the enlarged view the arrows point to spots where all three label co- localize. The scale bars correspond to 20 µm (overview) and 2 µm (enlarged picture). **C)** Four constructs that were used to develop intracellular labelling approach to visualize endogenous Ca_V_2.1 channels or manipulate their subcellular distribution. **D-F)** Left panels correspond to the expression of the intrabody-Halo-Tag-CCR5*, CRY2-* intrabody-Halo-Tag-CCR5 or CAG-Halo-Tag- intrabody-ΔCCR5 in Ca_V_2.1::neurons. Middel panels represent the expression of intrabody-Halo- Tag-CCR5, CRY2- intrabody-Halo-Tag-CCR5 or CAG-Halo-Tag-intrabody-ΔCCR5 in Cre-induced Ca_V_2.1-KO neurons by transfection of neurons with Cre-Td-Tomato. Right panel shows quantification of fluorescent intensities of CAG-Halo-Tag-intrabody-CCR5 (control: n_ROI_= 6, N_cultures_ = 2, Cre; n_ROI_ = 11, N_cultures_ = 2. Unpaired t-test, p-value<0.0001), CRY2- intrabody-Halo-Tag- CCR5 (control: n_ROI_= 10, N_cultures_ = 2, Cre; n_ROI_ = 10, N_cultures_ = 2. Unpaired t-test, p-value<0.0001), and CAG-Halo-Tag-intrabody-ΔCCR5 (control; n_ROI_ = 10, N_cultures_ = 2, Cre; n_ROI_ = 8, N_cultures_ = 2. Unpaired t-test, p-value = 0.412). Scale bar = 20 µm. **G)** Quantification of the colocalization of Ca_V_2.1 channels in bassoon-positive spots (0.60 ± 0.01, n_ROI_ = 23, N_cultures_ = 3), Ca_V_2.1 channels colocalize with Bassoon and Gephyrin (0.85 ± 0.02, n_ROI_ = 20, N_cultures_ = 3), and Ca_V_2.1 channels colocalize with Bassoon and Homer (0.94 ± 0.006, n_ROI_= 16, N_cultures_ = 3). **H)** Normalized intensity of Ca_V_2.1 channels in inhibitory synapses (1.00 ± 0.08, n_ROI_= 27, N_cultures_ = 3), and excitatory (1.08 ± 0.14, n_ROI_ = 30, N_cultures_ = 3) synapses, unpaired t-test, p-value = 0.612. Data are presented as mean ± SEM. **I)** Quantification of the occupancy rate of Ca_V_2.1-channels labelled with the halotag- intrabody against GFP and C-terminal anti-Ca_V_2.1 antibody (construct I: 0.47 ± 0.04, n_ROI_ = 11, N_cultures_ = 2, construct II: 0.75 ± 0.01, n_ROI_ = 30, N_cultures_ = 3, construct III: 0.82 ± 0.01, n_ROI_ = 29, N_cultures_ = 3). Data are presented as mean ± SEM. Number of replicates are summarized in suppl. table Fig.1.

Next, we added a photoactivatable cross-linker module based on cryptochrome 2 from *Arabidopsis thaliana* Cry2-olig (Taslimi et al., 2014) between the intrabody and HaloTag and found specificity of targeting was not affected (Fig.1 E, I). Probing the occupation of endogenous Ca_V_2.1 channels with the expressed intrabody confirmed that most intrabodies were linked to Ca_V_2.1 channels within the neuron (Fig.1I). In contrast to extracellular tagged Ca_V_2.1 channels (Watschinger et al., 2008), we found no difference between Halo- or GFP-tagged channels in current density and kinetics tested in heterologous expression.

Once verified, this approach to label endogenous Ca_V_2.1 channels was used to conduct fluorescent recovery after photo bleach (FRAP) experiments or single particle tracking (SPT) experiments to examine the dynamics of Ca_V_2.1 channels. In mature hippocampal neurons (15- 21 DIV), Ca_V_2.1 channels form distinct clusters in presynaptic boutons but can be also observed in dendrites and spines. Co-expression of the calcium sensor synaptophysin::jRGECO1a confirmed the co-localization of Ca_V_2.1 channels in presynaptic boutons (Fig. 2A-C). Small channel clusters outside synapses were frequently observed but their contribution to action potential (AP)-evoked calcium signals was negligible (Suppl. Fig. 2D). Importantly, calcium transients in synapses expressing intrabody-tagged Ca_V_2.1 channels were not affected as compared to control synapses not expressing the intrabody (Fig. 2F). Despite clear accumulation of Ca_V_2.1 channels in synapses, FRAP experiments revealed that on average 10-30% of the fluorescence recovers within five minutes after photo bleaching. However, the recovery rate strongly varied between individual synapses. Some synapses demonstrated fast recovery for up to 80% of baseline values in approximately five minutes, whereas in other synapses almost no recovery was observed within the same time interval (Suppl. Fig. 2A-C). These data demonstrated that synaptic population of endogenous Ca_V_2.1 channels is not static but can undergo local renewal within a few minutes. To explore this phaenomenon in more details, we employed SPT of individual Ca_V_2.1 channels to explore their spatiotemporal dynamics in functional synapses.

**Figure 2:**
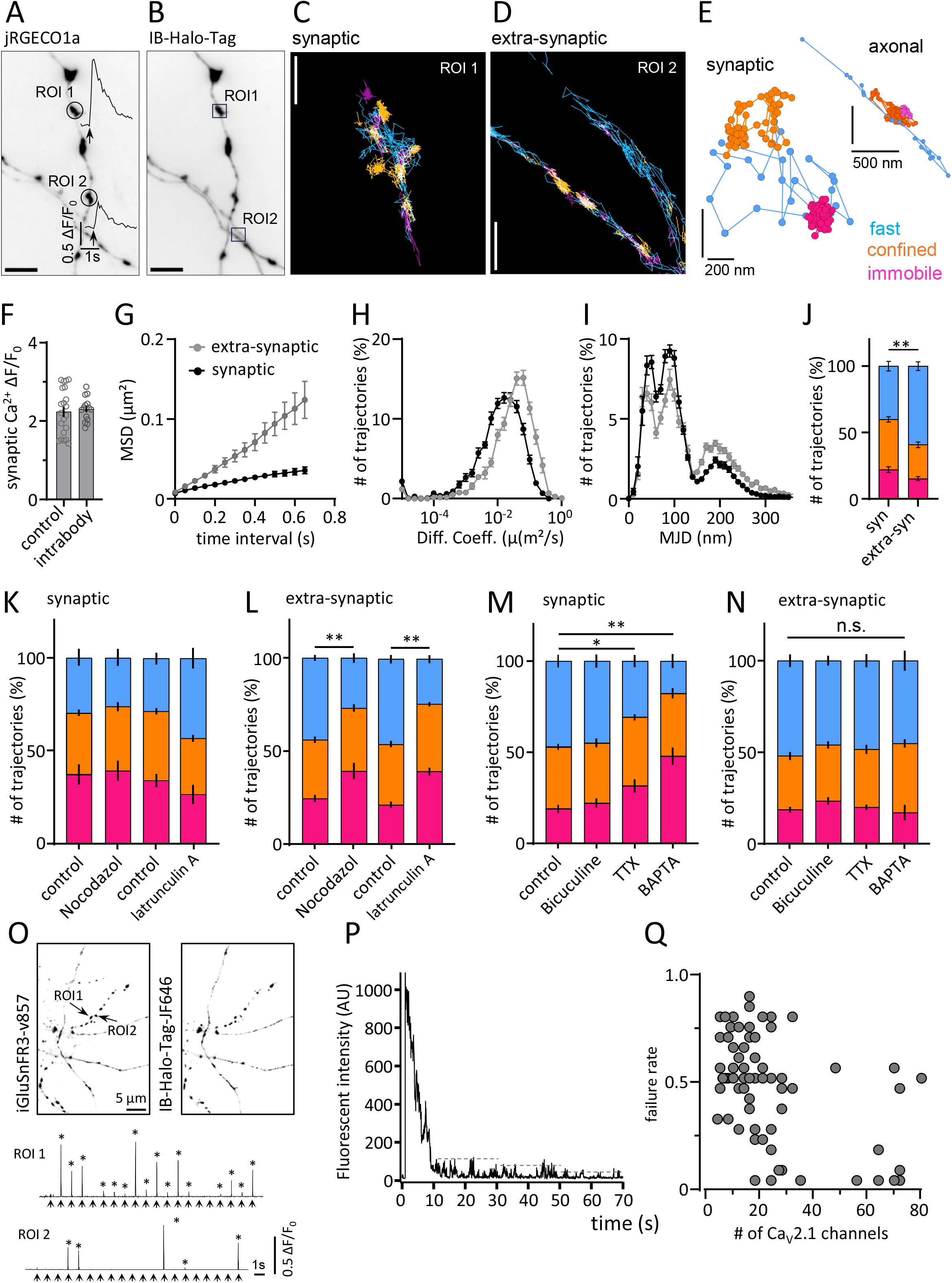
Dynamic of Ca_V_2.1 channels inside and outside synapses along the axon. **A)** Hippocampal neurons transfected with the calcium sensor jRGECO1a::synaptophysin, which also serves as synaptic marker. Calcium transients are shown for two regions along the axon. **B)** Same ROI as A) with expression of Halo-tag::intrabody, subregions are enlarged in **C)** and **D)** to illustrate individual trajectories of CaV2.1 channels inside a synapse and along the axon. **E)** Individual trajectories of synaptic and axonal localized Ca_V_2.1 channels, the colour code corresponds to immobile (pink), confined (orange) or free mobile (blue). **F)** Quantification of calcium transients from control neurons expressing jRGECO1a::synaptophysin (2.24 ± 0.12, n_synapse_ = 189, N = 3) and intrabody expressing neurons (2.30 ± 0.07, n_synapse_ = 135, N = 3) unpaired t-test, p-value = 0.707. **G)** Quantification of the mean square displacement (MSD) of Ca_V_2.1- channels in synaptic (black line) and extra-synaptic (gray line) compartments. **H)** Frequency distribution of the diffusion coefficient of Ca_V_2.1 channels in synaptic (0.027 ± 0.001, n_synapses_ = 205, N_cultures_ = 3) and extra-synaptic compartments (0.059 ± 0.004, n_axon segment_ = 184, N_cultures_ = 3), unpaired t test p-value<0.0001. **I)** Frequency distribution of weighted mean jumping distance (MJD) of Ca_V_2.1-channels in synaptic (black line) and extra-synaptic (grey line) compartments. **J)** Quantification of the proportions of different motions of Ca_V_2.1-channels based on MJD in synaptic and extra-synaptic compartments, Chi-square test, p-value = 0.026. Data are presented as mean ± SEM. **K-N)** Quantification of Ca_V_2.1-channel mobility in synaptic and extra-synaptic compartments based on MJD under different conditions. **K, L)** Preincubation of neurons with Nocodazol (1µM) or Latrunculin A (5µM) for 30 min. (Nocodazol: synaptic Chi-square test, p-value = 0.386; extra-synaptic Chi-square test, p-value = 0.001, Latrunculin: synaptic Chi-square test, p- value = 0.653; extra-synaptic Chi-square test, p-value = 0.001) **M)** Preincubation of neurons with TTX (1µM) for 30 min (synaptic Chi-square test, p-value = 0.0257; extra-synaptic Chi-square test, p-value = 0.911). **N)** Preincubation of neurons with BAPTA-AM (20 µM) for 30 min inside synapses (synaptic Chi-square test, p-value = 0.0063; extra-synaptic Chi-square test, p-value = 0.379). **O)** Example image of axons expressing iGluSnFR3 and intrabody::Halo-tag. Example traces of glutamate responses and bleaching curve (**P**) of synaptic concentrated Ca_V_2.1 channels are shown. **Q)** Correlation of failure rate with channel number of synapses, Pearson correlation coefficient r = -0.4637. Sample sizes for trajectories/synapses/cultures are given in Suppl. Table Fig. 2.

### Mobility of Cav2.1 channels differs within and outside synapses

To follow individual Ca_V_2.1 channels in synaptic and extrasynaptic locations, we used Halo-ligand Halo647JF (0.5 nM) (Fig. 2A-E). Tracking individual Ca_V_2.1 channels for up to 100 frames covering 5-s-long intervals revealed that a substantial fraction of axonal Ca_V_2.1 channels continuously move in and out of synapses. Moreover, SPT data showed high heterogeneity in mobility of Ca_V_2.1 channels when compared between synaptic and extrasynaptic locations. Inside synapses, we found a higher fraction of confined Ca_V_2.1 channels as compared to non-synaptic regions, where most channels move freely or exhibit directed motion (Fig. 2E).

Location-dependent calculation of the mean square displacement (MSD) and the distribution of the diffusion coefficients verified this clear difference in mobility between Ca_V_2.1 channels inside and outside synapses (Fig. 2 G-H). To directly assess the displacement per time interval, next we analyzed the mean jumping distance (MJD), which reflects an average distance covered by a channel within 50 ms, for trajectories longer than 3 time points (i.e., ≥ 200 ms). The weighted MJD for all trajectories revealed three distinct populations of channels in respect to their spatial displacement. We categorized MJD values shorter than 75 nm, within 75-150 nm range, and longer than 150 nm as immobile, mobile but confined and freely diffusive or actively transported Ca_V_2.1 channels, respectively. We found that a large population of endogenous Ca_V_2.1 channels (∼75%) is mobile within the synapse (Fig. 2H-J), confirming our previous data for overexpressed Ca_V_2.1 channels (Heck *et al*., 2019; Schneider *et al*., 2015). The proportions of immobile, confined and freely diffusive Ca_V_2.1 channels differ significantly between the synaptic and axonal (extra- synaptic) populations of Ca_V_2.1 channels (Fig. 2J).

Due to the intracellular position of the citrine tag, a subset of Ca_V_2.1 channels could potentially be detected not only in the cellular membrane but also in vesicles inside the axon or the presynaptic bouton. To examine this, we evaluated the calcium responses to electrical field stimulation along the axon expressing cytosolic GCaMP6f. We found that calcium signals were localized almost exclusively to presynaptic boutons co-labelled with synaptophysin::miRFP (Suppl. Fig. 2D-F). No calcium responses were evident for small channel clusters detected outside synapses, which confirmed intracellular localization of extra-synaptic Ca_V_2.1 channels. High mobility and directed motion of channels in the axon can be related to their occasionally observed active transport. To ascertain this, we interrupted active transport by incubating neurons for 30 min with either latrunculin A or nocodazole to depolymerize actin filaments or microtubules, respectively. Both treatments induced a clear reduction in the proportion of highly mobile Ca_V_2.1 channels outside synapses but did not affect mobility of synaptic Ca_V_2.1 channels (Fig. 2K, L).

Taken together, these results demonstrated that substantial fraction of axonal Ca_V_2.1 channels outside synapses are subject of trafficking and do not contribute to stimulation-triggered presynaptic calcium influx. In contrast, calcium imaging approach confirmed the membrane localization and functional relevance of synaptic Ca_V_2.1 channels. In the following experiments we focused on synaptic population of Ca_V_2.1 channels and addressed the question of how network activity and intracellular calcium concentration influence channel mobility.

### Suppression of the network activity decreases mobility of synaptic Ca_V_2.1 channels

To examine the impact of neuronal activity on channel mobility, we conducted SPT experiments under enhancement or suppression conditions of the network activity induced by blockade of GABA_A_ receptors (10 µM Bicuculine) or voltage gated sodium channels (1 µM TTX for 10 min), respectively. Treatment of neurons with TTX led to a significant increase of the confined and immobile populations of synaptic Ca_V_2.1 channels, whereas network disinhibition by Bicuculline did not affect Ca_V_2.1 channel mobility. Neither Bicuculline, nor TTX influenced the mobility of non-synaptic channels, hence additionally confirming lack of functional relevance of this channel population (Fig. 2M, N). These data corroborate our previous observations (Heck *et al*., 2019; Schneider *et al*., 2015) and shows a direct link between synaptic activity and mobility of synaptic Ca_V_2.1 channels.

To clarify whether the mobility of Ca_V_2.1 channels is calcium-dependent, we chelated free intracellular Ca^2+^ by incubating neurons for 30 min in 1 µM BAPTA-AM. We observed strong spatial stabilization of synaptic Ca_V_2.1 channels, while mobility of channels outside synapses remained unaffected (Fig. 2M-N). These data demonstrated that free intracellular calcium influences the mobility of Ca_V_2.1 channels. In addition, the results obtained with BAPTA treatment indicate that the well-known impact of BAPTA on the vesicular release probability is potentially also caused by the stabilization of Ca_V_2.1 channels in the synapse.

To directly test the impact of synaptic Ca_V_2.1 channels on the reliability of neurotransmitter release, we performed glutamate imaging using iGluSnFR3.v857 (Aggarwal et al., 2023). Glutamate release measurement allowed us to define the position of individual release sites and correlate it with topological distribution of Ca_V_2.1 channels within the same synapse. Synapses were activated at low stimulation frequency (1 Hz for 1 min) using a field electrode while postsynaptic glutamate receptors were blocked with CNQX and APV to suppress the network activity. To quantitatively assess the reliability of neurotransmitter release, the failure rate of evoked release events in response to repetitive stimulation was calculated. In parallel, the fluorescence of the tagged Ca_V_2.1 channels was used to calculate the number of channels per synapse. To achieve this, we used the bleaching steps of individual fluorophores to estimate the total number of calcium channels within the synapse based on 1:1 stoichiometry of channels to intrabody and HaloTag (Fig. 2O, P) (Ghelani *et al*., 2023; Hummert et al., 2021). We found that the Ca_V_2.1 channel numbers in individual synapses inversely correlate with the failure rate, confirming previous reports showing that abundance of Ca_V_2.1 channels in presynaptic membrane scales the release probability of the synapse (Holderith *et al*., 2012; Miki *et al*., 2017). However, the results of our FRAP experiments clearly demonstrate that Ca_V_2.1 numbers are not static but vary from synapse to synapse over time (Fig. 2 Q; Suppl. Fig. 2 A-C).

### Simulation of the Ca_V_2.1 distribution predicts relevance of mobile channels for synaptic reliability

The localisations of Ca_V_2.1 channels over time allows us to use segmentation tools to quantify the molecular organisation of channels along axons. Voronoï-based segmentation (Levet et al., 2015) indicated that Ca_V_2.1 channels preferentially cluster within synapses identified by evoked glutamate release (Fig. 3A-C). Within regions defined by the local density of localisations representing most synapses we could identify nanoclusters of Ca_V_2.1 channels based on an even higher density of localisations (Fig. 3D, E). Nanoclusters are seen in about 40% of synapses (Fig. 3A-C), comparable to the fraction of immobilised channels (Fig. 2I, J). The density of localisations within nanoclusters varied substantially, indicating different cluster size of Ca_V_2.1 channels (Fig. 3D, E). Interestingly many synapses did not show a distinct nanocluster, which implies a dominance of mobile but confined channels within these synapses.

**Figure 3.**
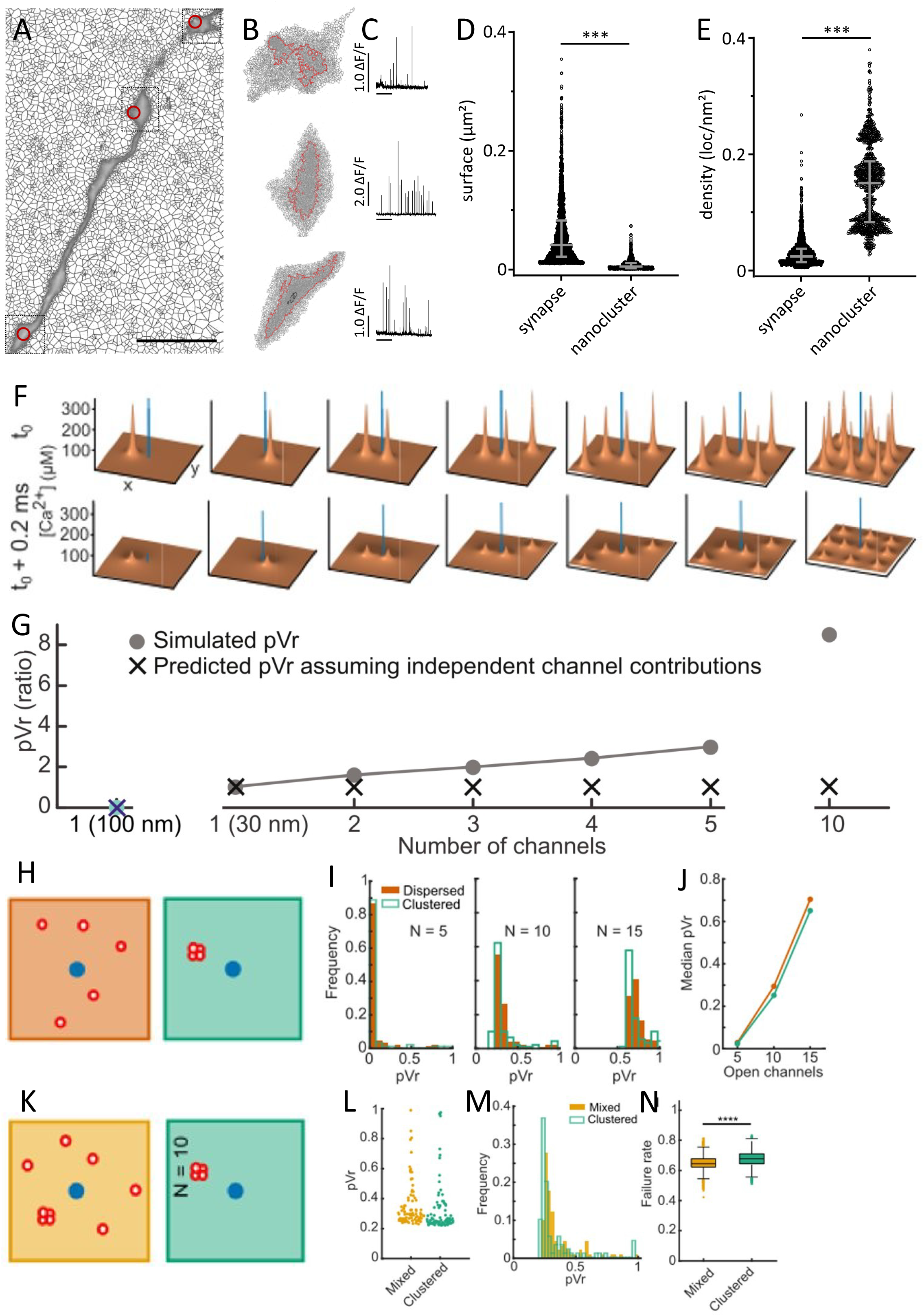
Characterization of vesicle release probability depending on location and number of open Ca^2+^ channels. **A)** Maximum projection of axonal segments expressing iGluSnFR3 stimulated 30x1Hz with a field electrode indicating several presynaptic locations. **B)** Voronoi segmentation of Ca_V_2.1 localisations recorded from SPT experiments. Tresholding based on the localisation density within neuronal structures allow to identify distinct regions of higher localisation density, often associated with synapses. Occasionally channels more clustered in nanodomains as exemplified in synapse 2 **(C)**. The example synapses are represented with their corresponding glutamate responses and density borders for synaptic objects and nanoclusters (black region within synapse 2). **D)** Surface distribution of objects and nanoclusters **E)** density of localisations in object and nanoclusters. Sample sizes for synapses/cultures are given in Suppl. Table Fig. 3. **F)** [Ca^2+^] spikes at time t_0_ of peak Ca^2+^ inflow (upper) and at a time of high vesicle fusion rate (t_0_ + 0.2 ms) (lower). Blue bar denotes the simulated release site location. Simulation area size is denoted by x = y = 0.5 µm. **G)** pVrs for each topology calculated as the ratio of each scenario compared to the pVr of the channel at 30 nm alone (second column). The grey line indicates the simulated pVr when the indicated channels were simulated together, whereas the black crosses show the pVr predicted assuming independent channel contributions, calculated from the individually simulated pVr of each channel (see Methods). (**H, K)** Schematic representation of the simulated channel arrangements. Dispersed (orange): channels placed independently at random positions. Clustered (green): all channels placed at the same randomly selected position. Mixed (yellow): half of the channels clustered and half dispersed.**I)** Distribution of simulated pVr values observed with N = 5, 10 and 15 open channels for dispersed and clustered conditions. **J)** Comparison of median pVr for N = 5, 10 and 15. (L + M) Distributions of pVrs for mixed and clustered conditions for N = 10 shown as scatter plot (G) and histogram (M). **N)** Distributions of simulated failure rates for mixed and clustered conditions for N = 10. Asterisks denote significant difference between mean values (****: p < 0.0001; two-sample t-test).

To quantitively assess the functional relevance of calcium channel number and spatial organization at a release site, we used mathematical modelling. We approximated the presynaptic active zone as a square surface (500 x 500 nm) and placed the vesicular release site in the center (blue bar in Fig. 3F). The parameters describing local calcium signals induced by a single AP-activated calcium were chosen to reproduce experimentally observed release probabilities for typical channel topologies (see Methods). We first evaluated the distance dependence of the vesicular release probability (pVr) by placing a single calcium channel either 100 nm or 30 nm away from the release site (Fig. 3F, G). With a channel 100 nm away produced virtually no release, a channel at 30 nm yielded a pVr of approximately 0.2, consistent with the established importance of tight coupling between calcium channels and release sites (Eggermann et al., 2012). The non-saturating pVr values predicted for single channels imply that additional channels could, in principle, contribute to release. However, because dispersed channels are broadly distributed and individual channel contributions decay steeply with distance, it is unclear whether distant channels contribute meaningfully to release. We therefore asked whether additional dispersed channels cooperate functionally despite their spatial separation. We investigated this by adding more channels at increasing distances and found that this caused a robust and steady increase in P_r_. Unexpectedly, the contribution of additional channels was largely independent of their spatial relationship to each other and instead depended primarily on their distance to the release site (Fig.3F, G, suppl Fig. 3) and even channels added 200 nm away from the release side significantly increased to PV_r_ (Fig.3F, G). Increasing the total number of channels within the active zone further confirmed both previous work and our simulations, showing a positive relationship between channel number and PVr (Fig. 2Q, 3G (Holderith *et al*., 2012; Miki *et al*., 2017). The effect was strongly cooperative, demonstrated by the observation that resulting total Pr was much higher than predicted by the combined individual effects of the single channels alone (Fig. 3G, see methods). These results indicate that dispersed channels cooperate functionally without the need for any direct molecular interaction while the spatial relationship between neighboring channels is of little relevance and their number and distance to the release site are major determinants of synaptic strength.

A key difference between clustered and dispersed channel arrangements is how they are affected by stochastic gating. When channels are clustered, stochastic outcome is partly compensated by combining several channels in one place, ensuring more reliable signals from that location. However, dispersion also increases the probability that at least one channel resides in close proximity to the release site, potentially improving release reliability. This, combined with the cooperative but independent effects additional channels have on Pr (see above), could be an advantage. We therefore tested this in simulations. If we simulate a dispersed versus a tightly coupled organization of channels, we see a slight shift in the distribution of P_r_ depending on channel number and organization. Few clustered channels (<8) show a similar P_r_ as dispersed channels, whereas >8 channels show a tendency to alter the distribution of the P_r_ (Fig.3H-J). The broader distribution of P_r_ values is clearly extended in direction of larger P_r_ values, meaning that the chance to trigger SV release in synapses depend on the local organization of channels and their dynamic within the membrane, reflected in a slight but significant reduction in the failure rate (Fig.3 K-N). Thus, channel dispersion is predicted to result in slightly more reliable synaptic output. We also investigated a mixed model of channel topologies where each half of the channels were clustered and dispersed, which resulted in more reliable responses than when all channels are clustered (Fig. 3K-N). If indeed local fluctuations of few channels are effective in tuning P_r_ values, we should see similar changes in case that we randomly cluster channels in the presynaptic membrane. A random clustering of channels we can address experimentally by fusion of CRY2-olig to the anti-GFP intrabody, allowing to optically cluster tagged Ca_V_2.1 channels as seen in overexpression experiments (Heck *et al*., 2019).

### Optogenetic clustering of Ca_V_2.1 channels

To directly control channel mobility and probe its contribution to neurotransmitter release, we introduced a light sensitive cross-linker (X-link) into the intrabody construct (Fig. 1C). Using a cryptochrome mutant (CRY2-olig) from *Arabidopsis thaliana* (Taslimi *et al*., 2014) enabled us to temporarily cluster Ca_V_2.1 channels within synapses. A single brief light flash (1 s/488 nm) induced a significant increase in the immobile and decrease in the mobile fraction of Ca_V_2.1 channels (Fig. 4D). The immobilization of Ca_V_2.1 channels did not affect the presynaptic calcium response or the number of Ca_V_2.1 channels in individual synapses (Fig. 4 C, E). However, evaluating the local density of Ca_V_2.1 channels before and after cross-link by Voronoï-based segmentation (Levet et al., 2015) revealed a significant decrease of Ca_V_2.1 channel nano-cluster size accompanied by an increase in the localization density (Fig. 4 F-I). Taken together, these experiments showed that light-induced clustering of Ca_V_2.1 channels does not lead to a recruitment of additional Ca_V_2.1 channels into the synapse but induces a substantial redistribution and compaction of pre-existing Ca_V_2.1 channels within the synapse.

**Figure 4:**
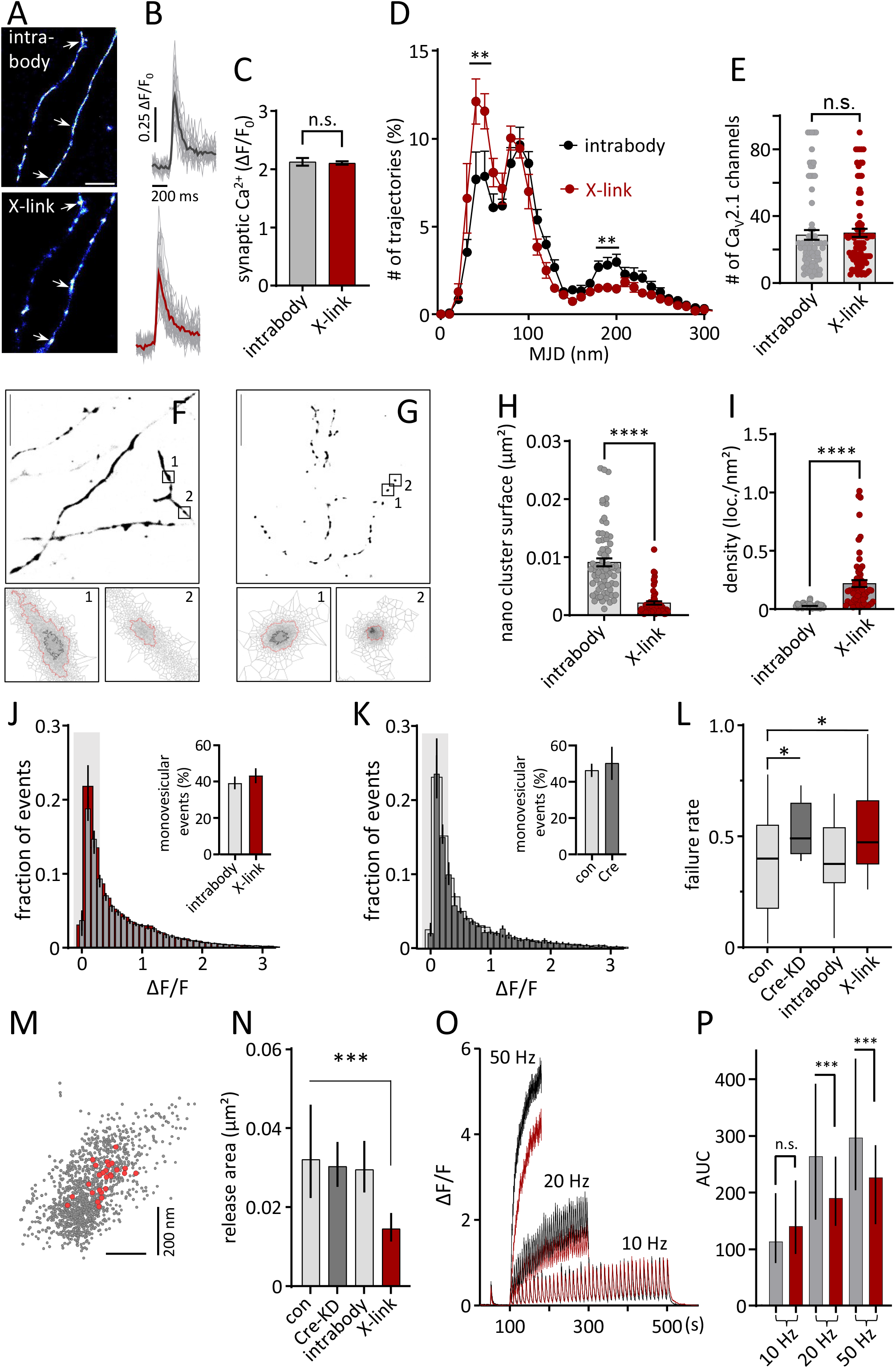
Organisation and function of Ca_V_2.1 channels before and after optogenetic manipulation of Ca_V_2.1 mobility. **A)** Examples of axons transfected with CRY2-intrabody-Halo-tag before and after blue light stimulation, arrows point to regions of induced clusters. **B)** Synaptic calcium transients of jRGECO1a::synaptophysin before and after blue light with average transient in gray (con) or red (CRY2). **C)** Quantification of synaptic calcium transients before light: ΔF/F_0_= 2.13 ± 0.07 and after light, ΔF/F_0_= 2.11 ± 0.03, unpaired t-test, p-value = 0.8. **D)** Frequency distribution of MJD of Ca_V_2.1 channels before (black line) and after blue light (red line). **E)** Quantification of synaptic Ca_V_2.1 channels number within synapses with and without x-link (Unpaired t-test, p-value = 0.735). **F, G)** Example images of Ca_V_2.1 localisations with and after light induced clustering. Accumulated Ca_V_2.1 localisations labelled with the intrabody::Halo-tag (F) or the intrabody-Cry2::Halo-tag. Selected regions (1, 2) correspond to presynaptic CaV2.1 channels distributed evenly or clustered within the synapse. The red line corresponds to the border of the synaptic localisation, black indicate the size of Ca_V_2.1 nano-clusters) **H)** Quantification of nanocluster area and **I)** localisation density before and after x-link in Ca_V_2.1 channels. **J)** Frequency distribution of glutamate responses to 1 Hz stimulation for 1 min from neurons co-transfected with intrabody and glutamate sensor. Fraction of monovesicular release events is not different between intrabody and x-link. **K)** Comparison between control and Cre induced KD reveal no difference of glutamate responses or fraction of monovesicular release. **L)** Failure rate of glutamate responses in the different conditions as indicated. **M)** Distribution of Ca_V_2.1 channels (gray dots) within a synaptic region overlayed by the localisations of glutamate release after 1 Hz stimulation for 1 min. **N)** Membrane area of glutamate release based on localisation of individual glutamate responses over time for the conditions as indicated. **O)** Average traces of synaptic glutamate responses to 51 stimuli of 10, 20 and 50 Hz. The immobilisation of Ca_V_2.1 channels (red) lead to a depression of transmitter release during repetitive stimulation in comparison to control conditions (grey). **P)** Area under the curve plotted for x-linked (red) and control (grey) conditions. Sample sizes for trajectories/synapses/cultures are given in Suppl. Table Fig. 4.

To evaluate whether the light-induced reorganization of Ca_V_2.1 channels affects the release properties of the synapse, we performed presynaptic glutamate imaging. The amplitude distribution of glutamate responses to a 1Hz stimulation train showed skewed but similar distributions under control condition, upon Cre-recombinase mediated KD of Ca_V_2.1 or when intrabody or X-linked intrabodies were expressed (Fig. 4J, K). Comparison between these distributions of evoked glutamate responses with spontaneous glutamate release in the presence of TTX (1 µM) (Suppl. Fig. 4A-G) confirmed that a substantial fraction (40-50%) of evoked glutamate responses is induced by multi-vesicular release, albeit this fraction was comparable across tested conditions (Fig.4 K, L). Next, we analyzed the response failure rate to evaluate the initial release probability of a synapse showing small changes in the simulation (Fig. 3I). Cre-induced downregulation of Ca_V_2.1 channels slightly increased failure rate (Fig. 4L), despite reported upregulation of Ca_V_2.2 channels (Cao and Tsien, 2010). This manipulation did not significantly change the amplitude distribution of glutamate responses or the fraction of mono- vesicular release events (Fig. 4K). An increase in the failure rate (Fig. 4L) indicated a specific function of Ca_V_2.1 channels for the reliability of transmitter release within glutamatergic hippocampal synapses. Interestingly, the X-link of Ca_V_2.1 channels also induced a small but significant increase in the failure rate (Fig. 4 L). These results suggest that redistribution of the mobile fraction of Ca_V_2.1 channels in the synapse indeed affect the association of channels with readily releasable vesicles in the presynaptic membrane.

Active zones exhibit multiple release sites that are distributed all over the presynaptic membrane area and can differ in their individual usage (Maschi and Klyachko, 2017; Mendonça et al., 2022). It is very likely that a dynamic redistribution of Ca_V_2.1 channels in the plane of the presynaptic membrane could influence either the release area, number of release sites or their individual usage over time, and underly the changes in failure rate observed in synapses after X-link. A deep learning-based algorithm Neuroimage Denoiser (Weißbach et al., 2024) applied to live glutamate imaging recordings allowed us to denoise acquired signals and substantially increase the signal-to-noise ratio (SNR). Using this approach, we utilized evoked glutamate responses to localize the position of individual release events over time in single synapses (Fig. 4M). In line with previous approaches using various fluorophores (Maschi and Klyachko, 2017), we identified 2-10 release sites per synapse spread over 0.025 ± 0.019 µm² area, which we designated as release area (Fig. 4N). The Ca_V_2.1 KD had no impact on the release area. Conversely, light-induced clustering of Ca_V_2.1 channels induced a substantial reduction in the spread of release sites within synapses (Fig. 4N), demonstrating that some release sites are not used after X-link. This is aligned with the observed compaction and lower mobility of Ca_V_2.1 upon light-induced clustering (Fig 4D, H, I).

A fraction of Ca_V_2.1 channels is often organized in nanoclusters as confirmed by Voronoï-based segmentation (Fig. 3A-E, 4F-I) but the use of release sites also depends on the local mobility of Ca_V_2.1 channels. This is potentially most critical by repetitive usage of release sites. To test this idea, we introduced repetitive field stimulations with 51 pulses delivered at 10, 20 or 50 Hz in neurons expressing the glutamate sensor iGluSnFR3. We calculated the area under the curve (AUC) of the induced fluorescence signals, because the used iGluSnFR3-sensor was too slow to resolve individual pulses at frequencies over 10 Hz but sensitive to follow changes in glutamate concentration. A single pulse preceding the train of stimuli was used to identify potential synaptic localizations along the axon and to distinguish spill-over responses from neighboring synapses. Comparison of responses to train stimulation between synapses expressing either the intrabody or the x-link construct revealed that transmitter release decreases at repetitive pulses above 10 Hz stimulation if channels are immobilized (Fig. 4O, P). This supports the idea that mobile Ca_V_2.1 channels enable alternative use of release sites in the presynaptic membrane and actively contribute to a reliable transmitter release.

In SPT experiments, we found that in control conditions Ca_V_2.1 channels are organized in nanoclusters (Fig. 3A-F, 4F) or are mobile but confined. Nanoclusters of Ca_V_2.1 channels are only detectable in a fraction of synapses (∼40 %), suggesting that synapses without Ca_V_2.1 nanoclusters might have clusters of other Ca_V_2 isoforms or have a looser organization of Ca_V_2.1 channels as suggested by FRAP experiments (Suppl. Fig. 2A-C). Quantitative analysis of synaptic population of calcium channel isoforms in cultured hippocampal neurons (18-24 DIV) showed that, in line with previous reports (Brockhaus et al., 2019; Cao and Tsien, 2010; Hoppa et al., 2012; Schneider *et al*., 2015), Ca_V_2.1 channels represent majority (60%) of the channel population, followed by Ca_V_2.2 channels (35%) (Suppl. Fig. 4H, I).

Rather moderate impact of the Ca_V_2.1 channel clustering on the failure rate and only effects on the average amount of released neurotransmitter at higher frequencies (Fig. 4K, L) raised a question of how relevant is a partially random distribution of synaptic Ca_V_2.1 channels in a glutamatergic synapse (only 40% contain Ca_V_2.1 clusters). Furthermore, the artificial nature of light-induced channel immobilization urged a search for endogenous mechanisms that control the available fraction of mobile Ca_V_2.1 channels.

Physiological changes in the release probability of a synapses are strongly influenced by the activation of metabotropic receptors, located in the pre- or postsynaptic membrane (Burke et al., 2018; Scheuber et al., 2004). The interaction and tight molecular association between GABA_B_Rs and Ca_V_2.2 channels (Muller et al., 2010; Schwenk et al., 2016) was proposed to be functionally important in homeostatic regulation of the synaptic release probability (Laviv et al., 2010). Nonetheless, we and others found that most calcium channels participating in neurotransmitter release in glutamatergic hippocampal synapses are Ca_V_2.1 channels, which are not tightly associated with GABA_B_Rs (Muller *et al*., 2010; Schwenk *et al*., 2016). Therefore, next we tested whether the heterogeneity in Ca_V_2.1 channel organization influences the consequences of GABA_B_R activation on the neurotransmitter release.

### Clustered Ca_V_2.1 channels are protected from G protein-mediated inhibition

Immunocytochemical experiments confirmed that 85% of presynaptic Bassoon clusters in hippocampal primary culture colocalize with GABA_B_R clusters in glutamatergic synapses, defined by the colocalization with Homer1 as postsynaptic marker (Suppl. Fig. 5). Activation of GABA_B_Rs, which are coupled to G_i_ protein, reduces presynaptic cAMP levels via the G protein α_i_-triggered pathway and mutes presynaptic calcium channels by liberating the β-γ subunits that directly interact with the pore-forming subunit of Ca_V_2.1 and Ca_V_2.2 (Herlitze et al., 1996). Previous studies have proposed that the heterogeneity of channel isoforms in the presynaptic membrane could serve as a mechanism to tune the impact of G protein-mediated modulation on transmitter release (Reid *et al*., 2003).

Within 10-15 min after activation of GABA_B_Rs with baclofen (5 µM), we observed a strong reduction of the presynaptic calcium responses measured by GCamp6::synaptophysin was evident (Fig. 5A, B). Calculation of the number of synapses that exhibited calcium responses prior to and after baclofen application showed that ca. 65% of synapses were fully inhibited upon GABA_B_R activation (Fig. 5C). Likewise, analysis of glutamate responses demonstrated that two thirds of synapses were quiescent after baclofen application. To rule out compensatory effects of Ca_V_2.2 channel upregulation in Ca_V_2.1 channel KD-neurons (Cao and Tsien, 2010; Schneider *et al*., 2015), we complemented treatment with baclofen by specific Ca_V_2.2 blocker conotoxin (1 µM). The fraction of synapses that remained active in presence of baclofen was not affected by acute block of Ca_V_2.2 channels in controls (Fig. 5C) but was strongly decreased (10%) in neurons with Ca_V_2.1 KD neurons. These results confirmed that a subset of synapses that show activity in presence of baclofen are equipped with Ca_V_2.1 channels, while Ca_V_2.2 channels dominate synapses that are fully inhibited by GABA_B_Rs (Fig. 5C). Proteomic studies have suggested only indirect coupling between Ca_V_2.1 and GABA_B_Rs (Muller *et al*., 2010; Schwenk *et al*., 2016) but it remains unknown whether the organization and mobility of Ca_V_2.1 channels can alter the GABA_B_R-mediated inhibition of release. We found that synapses remaining active after GABA_B_R activation were characterized by a much larger population of confined and immobile/clustered channels as compared to synapses that exhibited baclofen-induced suppression of release (Fig. 5D, E). This suggests that clustering or immobilization of Ca_V_2.1 channels can render synapses insensitive to GABA_B_R-induced inhibition. To test this hypothesis, we measured glutamate release after light-induced clustering of Ca_V_2.1 channels and baclofen application. Under control conditions, GABA_B_R activation dramatically increased the failure rate and the fraction of monovesicular release events (Fig. 5F-H). Light-induced clustering of the Ca_V_2.1 channels treated with baclofen also led to significant but less pronounced increase in the failure rate (Fig. 5G, H, J). No effect of X-link prior to baclofen treatment on the monovesicular release events was found, when compared to treatment with baclofen alone (Fig. 5G, J).

**Figure 5:**
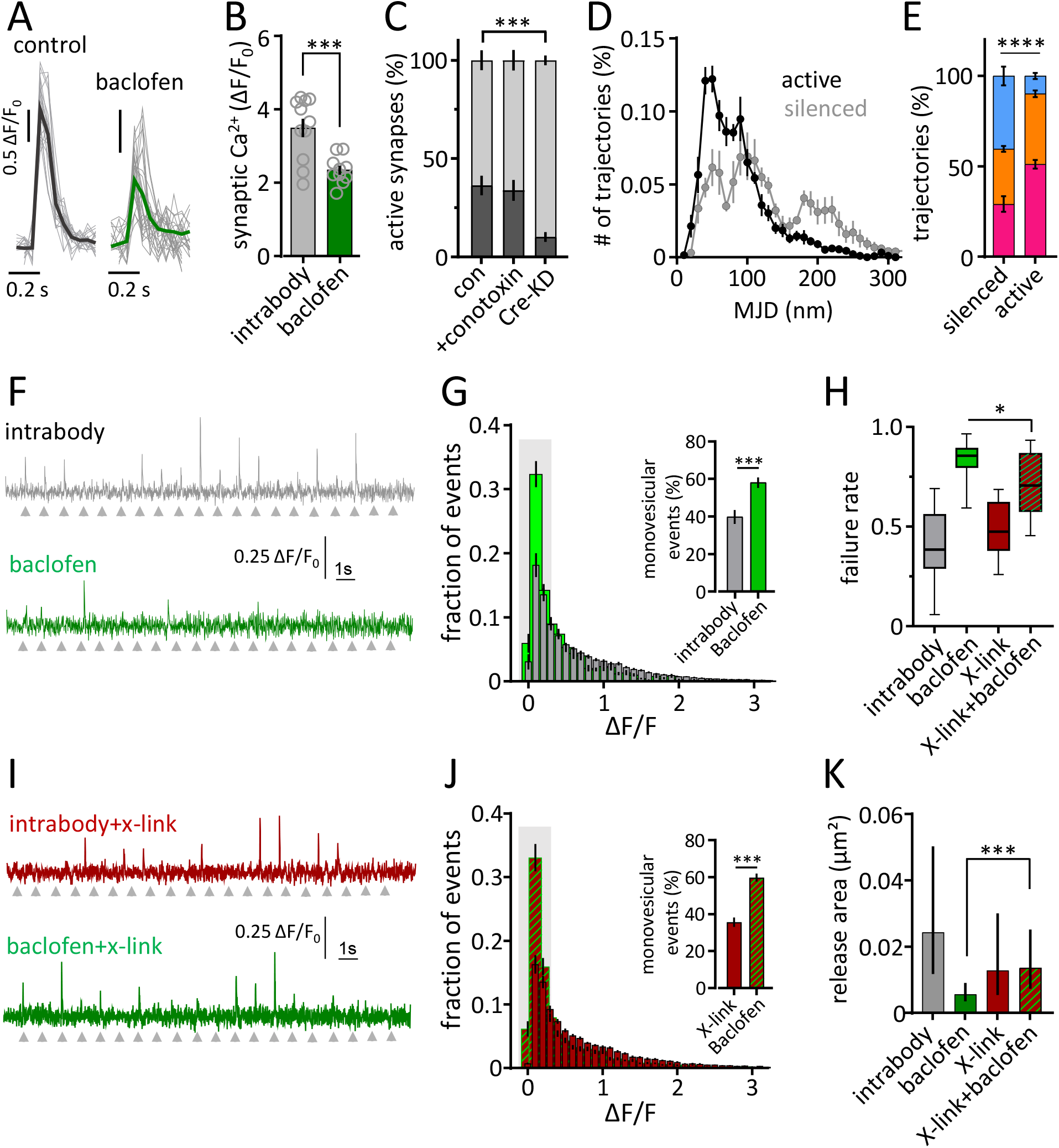
GABAB receptor mediated modulation of presynaptic release properties with and without manipulation of the Ca_V_2.1 channel dynamics inside synapses. **A)** Examples of presynaptic calcium transients in response to 1AP stimuli measured before and after application of baclofen (5 µM) using GCamp:synaptophysin. **B)** Quantification of calcium transients before and after baclofen application. **C)** Fraction of silenced (grey) and active (dark grey) synapses after activation of GABAB receptors for the indicated conditions. **D)** Distribution of mean jumping distance (MJD) for silenced (grey) or active (black) synapses after application of baclofen (5µM) **E)** Fractions of immobile (red), confined (orange) and fast moving (blue) Ca_V_2.1 channels in silenced and responding synapses (Chi-square test, p-value <0.0001). **F)** Example traces of glutamate responses with (green) and without (grey) activation of GABAB receptors or **I)** after x-link before (red) and after (green) activation of GABAB receptors. **G)** Frequency distribution of glutamate responses before (grey) and after activation (green) of GABAB receptors. The insert show fraction of monovesicular release under the different conditions. **J)** Similar data as in G) but after x-link of Ca_V_2.1. **H)** Quantification of failure rate for the indicated conditions. **K)** Release area calculated from the localisation of glutamate responses over time under the different conditions. Sample sizes for trajectories/synapses/cultures are given in Suppl. Table Fig. 5.

The evaluation of the localization of release events in the presynaptic membrane showed that only a small fraction of potential release sites was used. Light-induced clustering of Ca_V_2.1 channels reduced the number of release sites under control conditions but abolished a further reduction by baclofen (Fig. 5K). Thus, immobilization of Ca_V_2.1 channels partially counteracts the activity of GABA_B_Rs and provides additional evidence of the dynamic association of Ca_V_2.1 channels and release sites within the synapse as a potential mechanism to tune the reliability and robustness of synaptic transmission. Our results for metabotropic modulation of Ca_V_2.1 channel population demonstrate that it is dependent on the dynamic organization of channels. We found that the clustered Ca_V_2.1 channels are less sensitive, whereas dispersed Ca_V_2.1 and Ca_V_2.2 channels are more vulnerable to presynaptic GABAergic modulation. The mechanisms on Ca_V_2.1 channel stabilization within the synapse are manyfold. Extensively investigated earlier structural interaction between C-terminus of Ca_V_2.1 channels and presynaptic scaffold proteins affects synaptic targeting and retention of the channels (Chin and Kaeser, 2024; Held *et al*., 2020; Lubbert *et al*., 2017). Additionally, alternative splicing of the Ca_V_2.1 channel C-terminus was reported to affect the association and dynamics of Cav2.1 channels in the synapse (Heck *et al*., 2019) and can potentially tune the fraction of mobile versus clustered channels.

## Discussion

Most glutamatergic synapses in the cortex and hippocampus of the mammalian brain have only one single active zone facing a postsynaptic density (PSD) populated with few clusters of glutamate receptors (MacGillavry et al., 2013; Nair et al., 2013; Schikorski and Stevens, 1997). Despite these structural similarities, high variability in P_r_ of small synapses (Branco and Staras, 2009; Jensen et al., 2019) argues against a stereotypic molecular arrangement. One of the factors that support such variability in the hippocampus can be remarkably high synaptic turnover (Attardo et al., 2015; Bulovaite et al., 2022). In addition, high presynaptic variability can be considered rather as a benefit than noise with respect to information processing capacity and memory formation in recurrent neuronal networks (Kappel and Tetzlaff, 2024; Rusakov et al., 2020). We demonstrate here that the dynamic organization of presynaptic Ca_V_2.1 channels is another contributing factor that expands functional heterogeneity of synaptic population.

Local fluctuations of Ca_V_2.1 channels likely have no direct consequences for short-term plasticity within the first hundreds of milliseconds. Within this temporal window after the initial calcium influx, fast mechanisms alter the recruitment of SVs to the synaptic membrane and dominate acute short-term plasticity (Neher and Brose, 2018). However, at longer time scale the steady reorganisation of Ca_V_2.1 channels significantly contributes to reliability and robustness of release. Labelling of individual endogenous Ca_V_2.1 channels allowed us to monitor their dynamic distribution and assess the fraction of clustered, confined and freely moving Ca_V_2.1 channels (Fig. 2A-E). By manipulating the mobile fraction of Ca_V_2.1 channels via Cry2 mediated optogenetic clustering, we demonstrate that the mobile channel population increases SV release probability (Fig. 4L), defines the distribution and usage of release sites (Fig. 4M, N), enables transmitter release at higher frequencies (Fig. 4O, P) and is subject to GABA_B_R-mediated regulation (Fig. 5J, K).

Although, clustered Ca_V_2.1 channels were frequently found within synapses to serve canonical function of triggering the release (Chen et al., 2024; Indriati et al., 2013; Miki *et al*., 2017; Nakamura *et al*., 2015), several lines of evidence converge at the functional role of mobile Ca_V_2.1 channels. First, dramatic rearrangements within the presynaptic membrane during release and retrieval of SVs (Byczkowicz *et al*., 2018) raise the question whether Ca_V_2.1 channels can withstand such rearrangement and maintain their precise coupling to individual release sites. Second, distinct differences between Ca_V_2.1 channels and SVs in liquid-liquid phase separation-based organization support the flexible organisation observed in our experiments (Chin and Kaeser, 2024; Emperador-Melero *et al*., 2024). Third, the sensitivity of mobile Ca_V_2.1 channels to GABA_B_R-induced inhibition renders the possibility that metabotropic modulation can gradually manipulate the number of available Ca_V_2.1 channels, serve as gain control or high-pass filter (Burke *et al*., 2018) and additionally expand heterogeneity of synaptic P_r_. Fourth, alternative splicing of the distal C-terminus of the Ca_V_2.1 channels affects their affinity to presynaptic scaffolds and thereby the strength of coupling to release site (Heck *et al*., 2019; Lubbert *et al*., 2017). Fifth, alternative use of existing release sites increases the probability of release events at higher stimulus frequencies (Fig. 4O), hence supporting more reliable and consistent synaptic transmission. An additional aspect, which could be involved into the dynamics investigated in our study is a potential steady exchange of channels in the membrane. Exchange of Ca_V_2.2 channels has been suggested to require less than an hour (Cassidy et al., 2014), but turnover of Ca_V_2.1 requires further study accounting for potential difference between mobile and clustered channels in their residence time at the membrane.

Given dynamic organization of both pre- and postsynaptic sides, a flexible use of presynaptic release sites and postsynaptic receptor clusters seems beneficial and would favour persistent synaptic transmission. Inactivated VGCC can exchange with activatable VGCC and sustain the fraction of functional VGCC close to release sites. Our experimental data show that reliable use of release sites depends on the fraction of mobile Ca_V_2.1 channels. Unexpectedly, this functional cooperativity does not require physical interaction of channels. Instead, it is determined primarily by the distance of individual channels to the release site, while the spatial relationship between the channels have little influence. On average, synapses express more Ca_V_2.1 channels than Ca_V_2.2 and other channels, but large variability of FRAP in individual synapses (suppl. Fig. 4A-C) reflects the diversity of presynaptic Ca_V_2.1 channel population. Presence of both Ca_V_2.1 and Ca_V_2.2 channels with nearly identical kinetic properties (Li et al., 2007) in the same presynaptic bouton could be seen as redundancy. However, our experiments with GABA_B_R activation demonstrate that coexistence of Ca_V_2 channels and their organization at the presynaptic membrane reflects rather degeneracy and is instructive for reliable initiation of release in response to various patterns of presynaptic APs. Ca_V_2.2 channels are directly linked to GABA_B_Rs by KCTD proteins (Muller *et al*., 2010; Schwenk *et al*., 2016) and are the primary target of GABA_B_R- mediated inhibition. We found that synapses that remain active upon GABA_B_R activation have a larger fraction of stabilised/clustered Ca_V_2.1 channels (Fig. 5D) and are defined by a more robust nanostructure resistant to GABA_B_R-mediated modulation. There are several reports that Ca_V_2.1 channels differ in their response to G protein-mediated modulation (Reid *et al*., 2003; Scheuber *et al*., 2004). Taken together with the data on optogenetic clustering of Ca_V_2.1, these data demonstrate that mobile Ca_V_2.1 channels in hippocampal synapses can sustain neurotransmitter release at higher stimulation frequencies as compared to immobile/clustered Ca_V_2.1 channels, which however are more resistant to GABA_B_R-mediated inhibition. These mechanisms involving spatiotemporal dynamics of presynaptic Ca_V_2.1 channels provide a potential for presynaptic filtering and differential control of neurotransmitter release to dynamically modulate transfer of information in neuronal circuits. We propose that the presynaptic compartment imprint a rapid response code, which partially is depending on the mobile organisation of Ca_V_2.1 channels that tune the bandwidth of presynaptic release probability.

## Author contribution

A. El K., C. P. do A., C.W. and M.H. performed SPT-SMLM microscopy, glutamate and calcium imaging. A. El K. and J. H. optimized intrabody construct and optogenetic tools. M.D.M. and S.H. provided the knock-in mouse model and participate in discussion and data interpretation. A.B. and S.W. provided analyzing pipelines. A.S. and A.W. performed simulations. A. El K. and M.H. analyzed the data. M.H. provided funding and conceptualized the work, with input from A.W., J.H. and A.B.. M.H. wrote the initial manuscript, all authors contribute to discussion of final draft.

## Acknowledgment

We thank Anita Heine and Michela Borghi for preparation of neuronal cultures, as well as Nathalie Philipp and Michel Seiwert for their lab support. Further we thank Anna Bodzeta, Nasrin Sorusch and Junaid Akhtar for helping in initial experiments and fruitful discussions. We thank our students for participation in experiments, particularly Lucas Rüdiger, Patricia Przibylla, Kim Weidner and Tim Heidike. We thank the imaging core facility of the JGU and Christoph Rickert for support in imaging and data analysis. This work was supported by the Schramm foundation to M.H., the CRC1080, project B12 to M.H., the Novo Nordisk Foundation (Young Investigator Award, project ID NNF19OC0056047 to A.M.W.), and the European Research Council (ERC consolidator grant PlasticSite 101045054 to A.M.W.). The work was funded by the Deutsche Forschungsgemeinschaft (DFG; German Funding Foundation) MA 5806/7-1, project number 511099028 (M.D.M), MA 5806/1-2, project number AOBJ: 641682 (M.D.M), SFB1280 project number 316803389 (subproject A21, M.D.M) and GRK2862/1, project number 492434978 (subproject 05, M.D.M).

## Reagent and Resources

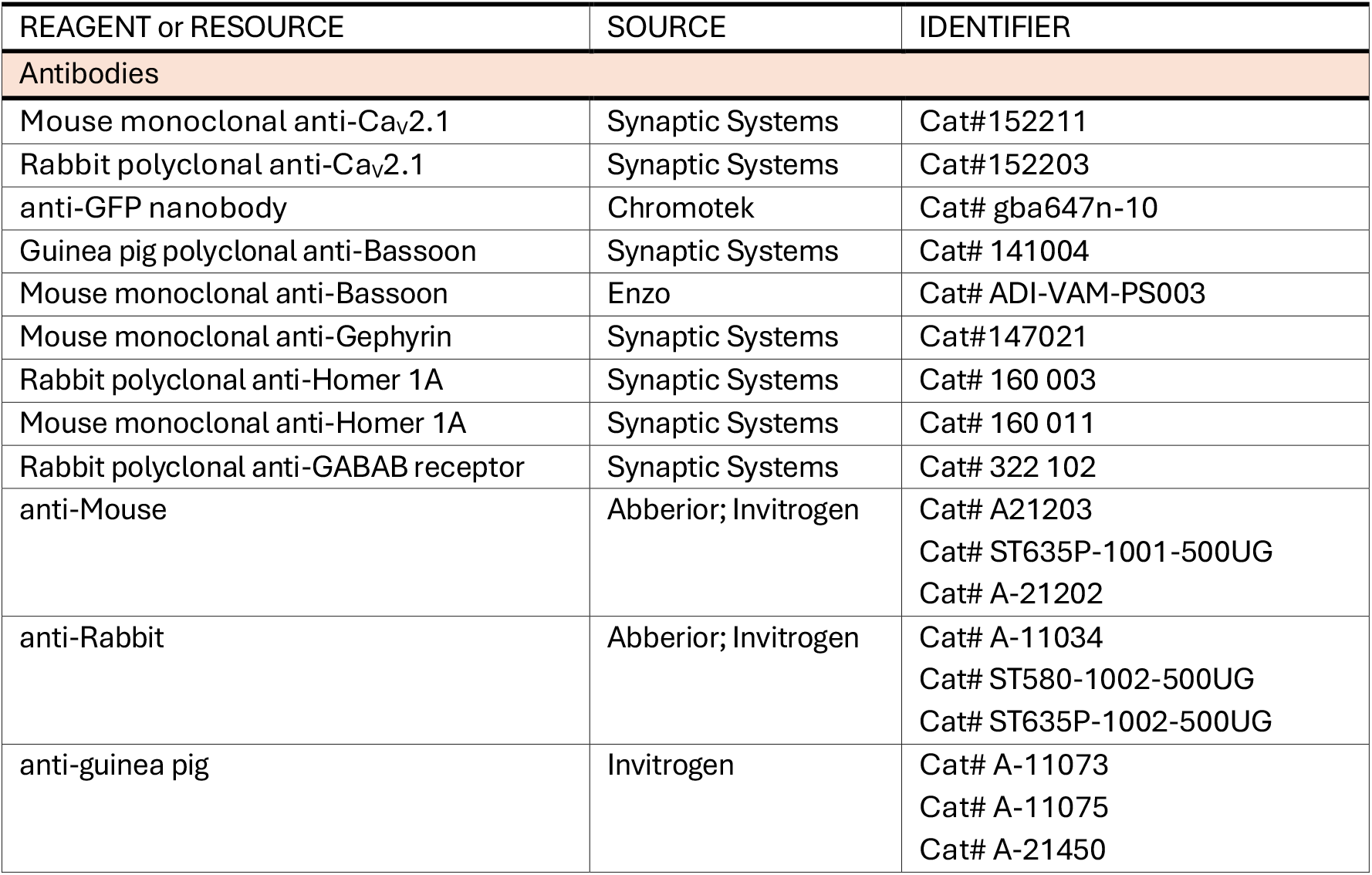
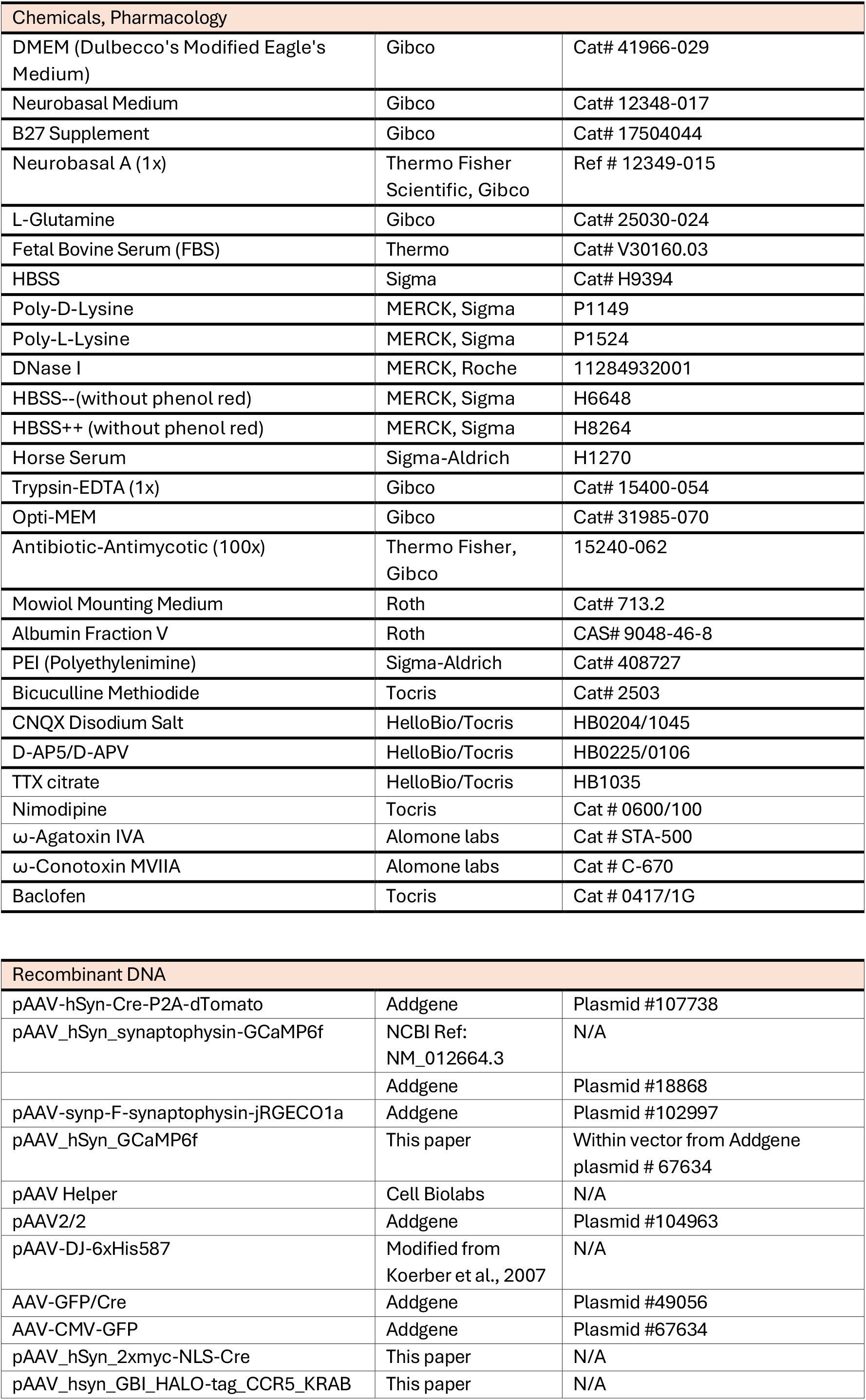
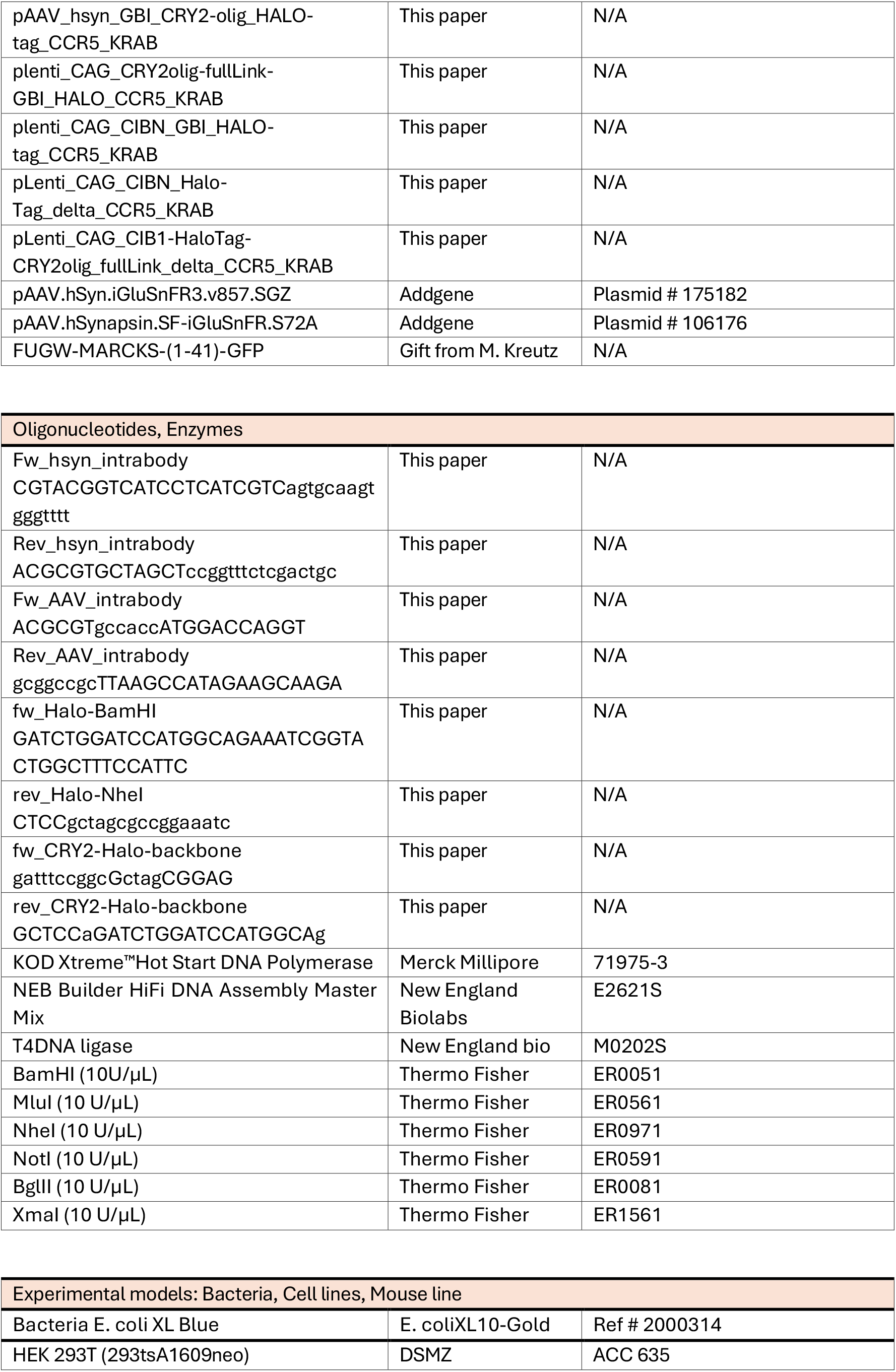
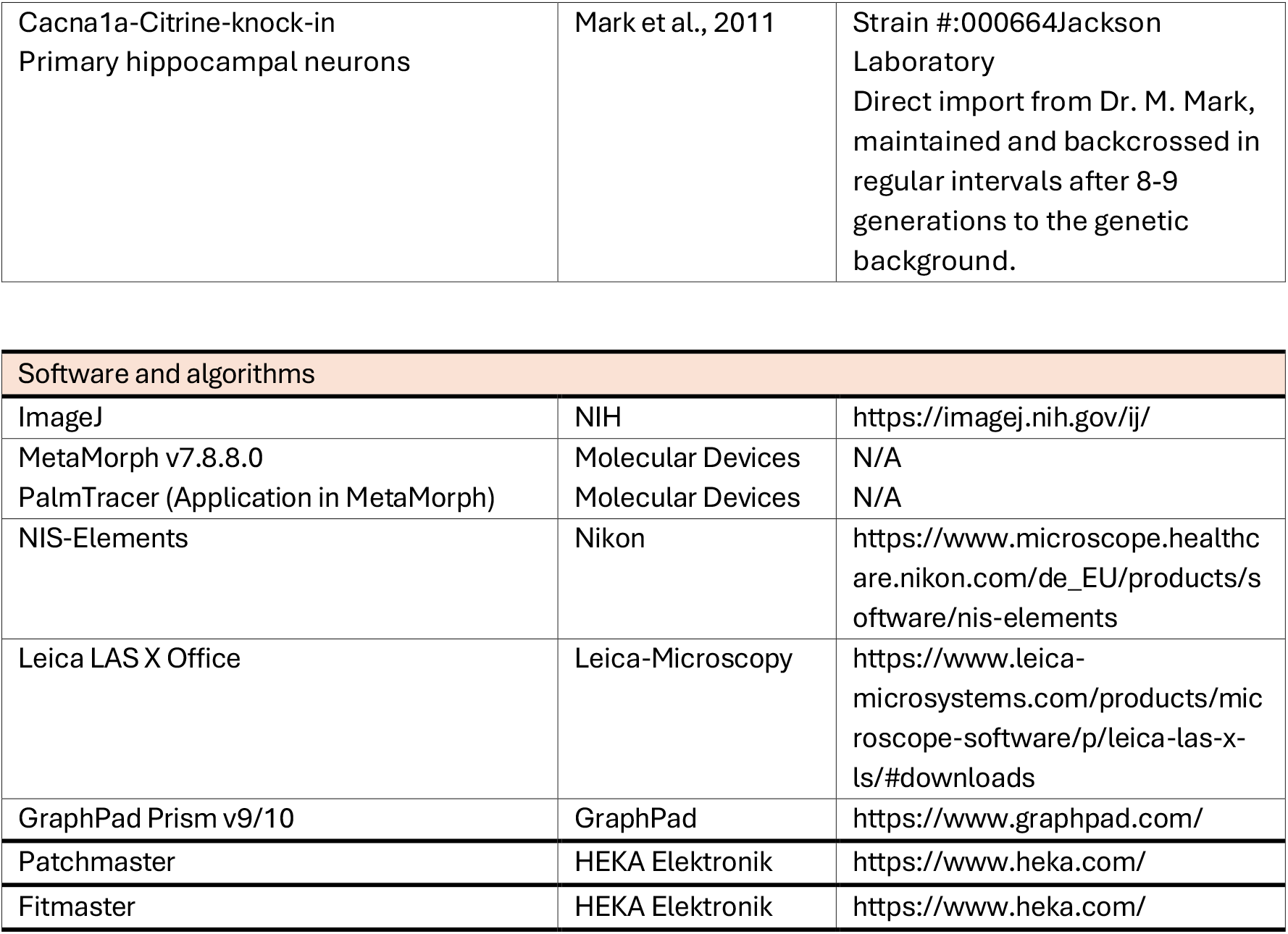

## Materials and Methods

### Cell culture and transfection methods

Primary hippocampal neuronal cultures were prepared as previously described (Beaudoin et al., 2012) from Cacna1a^citrine^ (Mark *et al*., 2011) newborn pups at postnatal days 0-1. Hippocampi were dissociated with trypsin and seeded in Dulbecco’s Modified Eagle (DMEM) medium supplemented with 10% fetal bovine serum and 1% L-glutamine. About 70,000 cells were seeded on 18 mm coverslips coated with poly-L-lysine. After 1h, the medium was replaced by Neurobasal A medium supplemented with 2% GlutaMax, 2% B27, and 0.1 M sodium pyruvate. Cells were incubated until use at 37°C in a humidified incubator with an atmosphere of 95% air and 5% CO_2_.

Human embryonic kidney (HEK) 293 cells were cultured in DMEM medium supplemented with 10% fetal bovine serum and 1% L-glutamine at 37°C in a humidified incubator with an atmosphere of 95% air and 5% CO_2_.

Primary hippocampal cells were transfected on DIV3-7 using a calcium phosphate transfection method (Jiang and Chen, 2006). In brief, for one 18 mm coverslip a mix of 24µl of ddH_2_O, 3µl of 2M CaCl_2,_ and 3µg of plasmid DNA was prepared and incubated for 5 minutes at room temperature. After incubation, 30 µl of 2xHBS solution (274 mM NaCl, 9.5 mM KCl, 1.4 mM Na_2_HPO_4_, 15 mM HEPES, 15 mM glucose, pH-7.14) were added and the mix was further incubated for 20 min at room temperature. During incubation, the neurons’ growth medium was replaced by pre-warmed Neurobasal A medium to which the DNA-calcium-phosphate mix was added. Cells were incubated for 40 min in the incubator to form the precipitate. After cells were washed two times with 1XHBS solution (pH 6.9) and once with Neurobasal medium. After washing steps, the original medium was returned to the cells.

HEK293 cells were transfected using the *polyethylenimine (PEI)* transfection method. For one well within a 12-well plate DNA and PEI were prepared in separated tubes. The ratio between cDNA and PEI was 1 to 3. Before mixing, 1 µg of Ca_v_2.1 α1_A_ subunit DNA, 0.5 µg of ß_3_ DNA, and 0.5 µg α_2_δ_1_ DNA were mixed in 48 µl of Opti-MEM. In a second tube 6 µl of PEI were mixed in 44µl of OptiMEM. Both solutions were incubated for 20 min at room temperature. After incubation, the cDNA and PEI solutions were fused and added to the cells. After 3h incubation the culture medium was replaced by DMEM medium supplemented with 10% fetal bovine serum and 1% L-glutamine. Cells showed robust expression of the channels after 48-72h.

### Immunocytochemistry

Hippocampal neurons were fixed in 4% PFA (37 °C) for 15 min and subsequently permeabilized with 0.3% Triton X-100/PBS for 10 min at room temperature (RT). Cells were washed three times for 10 min at RT using a washing buffer of the following composition: 1 x PBS; 2% BSA; 25 mM glycine, 1/10 of 10X Casein. Next, neurons were incubated with primary antibodies for up to 3 h at RT. After three times wash (10 min), the fluorescent conjugated secondary antibodies were used depending on the species of primary antibodies. After three washing steps using washing/blocking buffer, cells were mounted in Mowiol.

The specificity of antibodies against Ca_V_2.1 channel epitopes were probed in hippocampal neurons after Cre-recombinase induced knock-down of Ca_V_2.1 channels. Infection of hippocampal neuronal culture from floxed Cacna1a^Citrine^ mouse line took place on DIV 3 by use of a AAV8 virus caring Cre-recombinase::mCherry. Due to the delayed expression of Ca_V_2.1 the Cre-dependent knock down efficiency was close to 100%, which let allow to probe how specific available antibodies against Ca_V_2.1 channels are. We tested a monoclonal mouse anti-Ca_V_2.1 antibody (synaptic system, cat#152211) as well as a polyclonal rabbit anti-Ca_V_2.1 antibody (synaptic system, cat#152203) targeting the C-terminus of Ca_V_2.1 channels (Synaptic systems, Göttingen). As secondary antibodies we used donkey anti-mouse IgG Alexa Fluor 488 (Invitrogen Invitrogen; Cat#A21202) or goat anti-rabbit IgG Alexa Fluor 488 (Invitrogen; Cat# 11034).

Synaptic co-localization of Ca_V_2.1 channels in hippocampal cultures 17-21 DIV was probed by labelling the presynaptic scaffold protein Bassoon and the postsynaptic scaffold Homer 1A or Gephyrin to identify excitatory and inhibitory synapses, respectively. Images were acquired using a confocal microscope (Stellaris, Leica). The specificity of the Halo-tagged intrabody construct was assessed by incubating the transfected neurons with the Halo-ligand JF 646 (0.5 nM) for 1-3 hours prior fixation. After fixation and permeabilization the anti-Ca_V_2.1 antibodies (1/500) against the C-terminus were used to co-label Ca_V_2.1 channels on the C-terminus. The degree of co-staining between the Halo-tagged intrabody and the C-terminal antibody was taken as measure for the occupation of the Ca_V_2.1 channels with the intrabody construct.

The function of the transcription feedback loop was tested by expressing the intrabody-Halo-tag- CCR5 or the intrabody-halo tag without feedback control (ΔCCR5) in neurons prior infection with AAV8_hsyn-mCherry-Cre virus to knock-down Ca_V_2.1 channels and probing the distribution of intrabodies. The distribution of intrabodies was assessed in neurons 14 DIV after transduction. After Halo-Tag labelling with JF646 (1h with 5 nM ligand), probes were fixed and evaluated using a confocal microscope (Stellaris, Leica).

### Confocal image acquisition and colocalization

Confocal images were acquired using a Leica Stellaris 8 microscope with the Leica LAS X Software (Version 4.5.0.25531). The used microscope was equipped with an external light source (Leica EL6000) and a with light laser. Images were acquired at 400 Hz scanning rate using a 100x / 1.4 NA oil immersion objective (HC PL APO CS2) and unidirectional scanning of 1024x1024 pixels with a pixel size of 38.7 nm. Typically, images in x-y were acquired with a step size of 120 nm in z-direction.

Synaptic co-localization of Cav2.1 channels and synaptic markers was performed on maximum projections of Z stacks after background subtraction in ImageJ (rolling ball, 50 px). The quantification of colocalization was based on the SODA algorithm (Statistical Object Distance Analysis) within the ICY software (Open-source software: https://icy.bioimageanalysis.org/; (Lagache et al., 2018)). Spot detection in ICY was used to generate binary images for cluster analysis. We used a scale 3 threshold to optimally detect the signal with the size of 7-13 px diameter, and the center of detected spots was used as their spatial position. Then, we proceeded with the colocalization studio plug-in using a radius of 8 pixels (pixel size: 38.7 nm) that was used to automatically quantify the spatial coupling between different channels. The percentage of co-localisation between spots, cluster number, intensity and area were extracted and plotted using Graph-Pad Prism software (version 10).

### Recombinant DNA and Cloning

The pLenti-CAG-CIB1-Halo-Tag-intrabody construct serves as an anti-GFP intrabody targeting GFP or its derivative such as the citrine present in our Cacna1a^Citrine^-KI mouse model. The original plasmid pCGA-CIB1-GBI-mNeptune-CCR5/KRAB(A) was provided by Dr. Matthieu Sainlos (Bordeaux) containing the GFP binding protein (GBP) referred further as GFP-intrabody. Here we replaced the mNeptune with a HaloTag7 (Addgene #64691, doi: 10.1038/nchem.2002.). The HaloTag7 sequence was amplified from TUBB5-Halo. A unique restriction site BamHI was inserted at the N-terminal of the HaloTag7 sequence and the C-terminal of intrabody anti-GFP sequence using primers 5 and 6 (see table), respectively. The Halo-Tag fragment was inserted into the target backbone vector via the unique restriction site BamHI.

For cross-linking constructs via Cryptochrome, the CIB1 domain fragment was replaced by CRY2- olig isolated from the plasmid Addgene # 60032; (Taslimi *et al*., 2014). CRY2-olig was amplified by PCR with insertion of Xmal restriction site in the N-terminal of CRY2-olig sequence together with 20 nucleotides overhanging with the target vector (primer 9), and a C-terminal 20 nucleotides complementary overlap to the linker region of the intrabody (primer 10).

In a second PCR, the linker-intrabody carrying region of the pCGA-CIB1-GBP-mEOS- CCR5/KRAB(A) plasmid was amplified having the N-terminal 19 nucleotides (primer 11) complementary overlapping to CRY2-olig and a N-terminal BgIII restriction site with 26 nucleotides (primer 11) overlap to target vector. Both fragments were fused by a third PCR, then the resulting fragment was inserted in the target vector via the unique site Xmal and BgIII. After generating pLenti-CAG-mEOS-intrabody-CRY2-olig, the mEOS was replaced by the Halo-Tag via restriction enzymes specific to the restriction sites Pas1 and Nhe1, the Halo-Tag sequence was amplified by PCR from TUBB5-Halo (Addgene plasmid # 64691, (Uno et al., 2014)), having the N-terminal 23 nucleotides containing a unique restriction site BamHI (primer 8), complementary overhanging with the Halo-Tag and a C-terminal 25 nucleotides complementary to the intrabody sequence (primer 7). The Halo-Tag fragment was inserted into the target vector via the unique restriction site BamHI.

For the pAAV-hsyn-intrabody-Halo-Tag-CCR5/KRAB(A) construct, the whole backbone from pLenti-CAG-intrabody-halotag-CCR5/KRAB(A) plasmid except the CIB1 fragment was amplified by PCR (primers used: 3 and 4). The human synapsin promoter (hsyn) from the original plasmid pAAV2_hSyn_SP-Halo-STIM1 using oligonucleotides containing the Zinc-finger sequence at the upstream of hsyn sequence (primers used: 1 and 2), restriction sites specific for NotI and MluI. The fragments hsyn and intrabody-halotag_CCR5-KRAB(A), were ligated using the unique sites of NotI and MluI.

### Functional imaging of pre-synaptic calcium and glutamate responses

Live imaging experiments were conducted on mouse hippocampal cultures 18-23 DIV. All experiments were carried out at physiological temperature between 33-37°C. Neurons were perfused with extracellular solution containing (in mM): 145 NaCl, 2.5 KCl, 10 HEPES, and 10 D-glucose 2 MgCl_2_, 2 CaCl_2_ (pH 7.4, 300-310 mosmol). To suppress network activity and isolate presynaptic responses we blocked NMDA and AMPA receptors by adding D-AP5 (10 µM) and CNXQ (10 µM) to the extracellular solution. For image acquisition we used a Nikon Eclipse-Ti2-E inverted microscope equipped with an oil-immersion TIRF objective (60X, NA 1.49). Using the magnification switch of 1.5 x resulted in an effective pixel size of 72x72 nm. Images were acquired with an sCMOS camera (Orca-fusion C14440-20UP from Hamamatsu) under the control of NIS-Elements Advanced Research acquisition software (Nikon). Used fluorophores were excited using different laser lines (488/561 and 646 nm). For single particle tracking we used an oblique illumination mode to reduce background from out of focus fluorescent ligth. The acquisition rate for single molecule tracking or calcium imaging was 50 ms (20 Hz), glutamate signals were acquired with a frame rate of 200 Hz for iGluSnFR-S72A and 100 Hz for iGluSnFR3 within subregions of 512x512 or 700 x 700 pixels. Synaptic responses were evoked by electrical field stimulation using two platin electrodes placed within the recording chamber (Ludin chamber type1, LIFE IMAGING SERVICES, Basel Swiss) placed with ∼10 mm distance on the coverslip. Applied stimuli were 0.7-1 ms long with initial intensity of 50 mA to trigger action potential like responses. Trigger sequences were generated by the use of a stimulus generator (ACM Systems, model 2100) connected to an isolation unit (World Precision Instruments, #A385).

Glutamate or calcium sensors (iGluSnFR, jRGECO1a::synaptophysin) with or without co-expression of the intrabody was induced at DIV3-5 using the calcium phosphate transfection method. Neurons were used at DIV 15-20 after transfection. Calcium transients were recorded in response to single or repetitive stimuli of 3-10 pulses at 50 Hz. To induce optogenetic clustering of Ca_V_2.1 channels neurons were transfected with the CAG-HaloTag-intrabody-CRY2-olig construct 3-5 DIV and imaged after 15-20 DIV. Cross-linking was induced by a brief exposure to blue light (488 nm, 5% of initial laser power) for max 0.5 seconds. The clustering effect was evaluated immediately after light induced cross-link. The single light pulse induced clustering was effective for up to 10 min (Heck *et al*., 2019). Activation of GABAB receptors to access the metabotropic modulation of mobile Ca_V_2.1 channels was induced by preincubation of the neurons with 5 µM Baclofen (Tocris, UK) for 10 min, before probing presynaptic calcium or glutamate responses. The average quantal size of the glutamate response was accessed by blocking sodium channels with 1 µM TTX (Tocris, UK) 5 min before recording.

### Image analysis and quantification

Calcium transients were quantified after background noise was subtracted from image stacks using the rolling ball background subtraction in ImageJ. Hippocampal neurons expressing jRGECO1a::synaptophysin showed defined localisation of the calcium signals in synaptic boutons, in opposite to postsynaptic scaffolds of glutamatergic or GABAergic synapses. Detection of synaptic regions was performed by manual selection of synaptic puncta within the maximum projection of image sequences. The analysis of the calcium transients was performed using a custom-written macro in Image J software (Fiji). Synaptic calcium transients exceeding a threshold of 2.5. fold standard deviation (SD) of the averaged background fluorescence were accepted for further analysis. The baseline fluorescence (F_0_) was calculated from 20 frames before stimulation for pre-pulse normalisation of evoked signals within the selected regions. Calculated values for fluorescent changes (ΔF/F_0_) under different conditions were evaluated within Graph Pad Prism (version 10) to define differences in the relative amplitude changes.

Glutamate signals were extracted after denoising the image stack based on a deep learning algorithm (Weißbach *et al*., 2024) followed by background subtraction in ImageJ. Repetitive measurements of glutamate responses from individual synapses were possible using the iGluSnFR3-V857 glutamate sensor (Aggarwal *et al*., 2023). This sensor is characterized by its high affinity to glutamate (∼8.2 ± 0.1 µM), improved brightness, and photostability (Holderith *et al*., 2012). This allowed to monitor synaptic responses, failure rate, and quantal size of glutamate release from individual presynaptic terminals for up to 100 s without substantial loss of response amplitude. To monitor the initial release probability of synapses we used 1 Hz stimulation protocol. Defining individual glutamate responses was done by using a custom-written macro in Python (https://github.com/andreasmz/neurotorch). Signals were extracted from ROIs with 6 pixels radius. Individual transients were detected within a defined window of 1-3 frames after the stimulus frame. The fluorescent change was calculated as ratio between the baseline fluorescence (F_0_) calculated from 20 frames before stimulation and the F_max_ 1-3 frames after the stimulus evoked signal within the selected regions. Detected responses were extracted only from axonal segments. The quantal size for glutamate responses was extracted from recordings in the presence of 1 µM TTX, which occurred stochastically along axons and excided 3 SD of the background noise. Co-transfection with HaloTag-intrabodies or jRGECO1a::synaptophysin confirmed that detected signals correspond to clusters of Ca_V_2.1 channels or synaptic vesicles along axons and do not present crosstalk signals from crossing axons or dendrites within the network. Frequency distribution and failure rate were quantified and compared using Graph Pad Prism (version 10).

### Localisation of release sides and release area

Glutamate signals were used in addition to define the location of preferred release sites within the synapse. As seen in Fig. 4J, K; suppl.Fig.4C-G, a certain fraction of responses shows multivesicular release events, which indicate that we will not be able to define single release sites. But the individual responses over time could be used to define the release area within the presynaptic bouton. For this analysis individual synapses were cropped out of denoised and background subtracted images and used to better define the localisation of glutamate events using the ImageJ plug-in ThunderSTORM (Ovesný et al., 2014). Calculating local maxima within a radius of 7 pixel and adjusting the peak intensity threshold allowed us to define the subpixel localisation of individual release sites based on gaussian fitting of the fluorescent intensity profile. For the localisation we used the maximum response within the first three frames after the stimulus. Individual events where localised over time and replotted within the ROI, converged into a polygone selection and fitted by the convex hull plug-in in ImageJ to define the surface of the release area between the localized release sites of individual synapses. Distribution and statistical analysis of release site area under different conditions were analysed in Graph-Pad Prism (version 10).

### Fluorescent recovery after photo bleach (FRAP) experiments

Hippocampal neurons transfected with jRGECO1a::synaptophysin and the HaloTag-intrabody were used for FRAP experiments to localise presynaptic boutons and define the local turnover rate of Ca_V_2.1 channels within the photobleached area. We used a 2D Galvo-Scan System coupled to the TIRF-microscope (Nikon) to select identified synaptic regions and bleach diffraction limited spots within the field of view using a 60x 1.49 NA objective. Recovery of the fluorescent was immediately monitored at 5 Hz acquisition rate for the first minute and 1 Hz acquisition rate for the following time until recovery reached a steady state, usually within 4-5 min after bleaching. The recovery rate was normalized to the average fluorescent intensity of ten frames before photo bleach, which was quantified in GraphPad Prism (version 10).

### Single-particle tracking microscopy (SPT)

SPT experiments were performed on mouse hippocampal cultures 15-23 DIV at 37°C. Neurons were perfused with extracellular solution containing (in mM): 145 NaCl, 2.5 KCl, 10 HEPES, and 10 D-glucose 2 MgCl2, 2 CaCl2 (pH 7.4). Experiments were conducted with a Nikon Eclipse-Ti2-E inverted microscope using oblique illumination to reduce background fluorescence and define single fluorophores. The acquisition rate was 20 Hz with up to 5000 frames in ROIs 512x512 pixel. The focal plane was stabilized by using the perfect focus system from Nikon for Z focus stabilization. In all experiments we used the Halo-ligand 646JF prediluted at a concentration of 200 µM in DMSO. Neurons were labelled by adding the ligand to the culture media for 1-2 h at a final concentration of 0.5 nM. The fluorophore was excited using a 640 nm laser. Crosslinking of Cry2-olig labelled Ca_V_2.1 channels was induced by a brief illumination with a 488 nm laser line.

Localisation and tracking of individual fluorophores was performed using either PalmTracer software plug-in for MetaMorph (Izeddin et al., 2012) or using ThunderSTORM plug-in in ImageJ followed by analysis of trajectories and jumping distance using Swift (custom software, Endersfelder Uni Bonn). Trajectories of single Ca_V_2.1 channels were reconnected assuming a maximum displacement of 3 pixels (pixel size of 72 nm) based on visual inspection of the movies. Spots were detected by thresholding the images and localized by fitting a 2D Gaussian function with a 9-pixel radius that matches the microscope’s point spread function and an initial sigma of 1.6, achieving ∼35 nm localization accuracy in both x and y dimensions. Based on trajectories longer than 8 time points average diffusion coefficient was calculated based on the mean square displacement (MSD) versus Δt curve slope (D = MSD/4Δt) for the first 4 points. Weighted jumping distances were calculated based on trajectories with a minimal length of four time points. The localisation of trajectories within synapses or along axons was based on diffraction limited images of jRGECO1a::synaptophysin. Based on the frequency distribution of weighted jumping distances the mobility of Ca_V_2.1 channels was classified in three populations, immobile (average displacement ≤ 75 nm), confined (average displacement 75 – 150 nm) and mobile (average displacement > 150 nm). This classification was confirmed by averaged MSD plots for respective trajectories. Statistical analysis and graphical illustrations were performed using GraphPad Prism (version 10).

### Calculation of channel number and local channel density

Number of Ca_V_2.1 channels within synapses were analysed based on photo bleaching over time. The confined localisation of the majority of synaptic Ca_V_2.1 channels was bleached after prolonged excitation of the HaloTag-intrabody labelled channel population. The fluorescent signal was completely bleached within less than 5000 frames (20 Hz acquisition rate), which allowed identification of single fluorophores as one-step photobleaching events (see Fig. 2P). Recordings were background subtracted and synaptic regions chosen (10 pixel diameter) to identify single bleaching steps at the end of the bleaching curves. The used fluorophore JF646 showed sufficient photostability under physiological conditions to identify individual bleaching steps. For the photo bleaching step analysis of individual synapses we used the algorithm introduced by (Hummert *et al*., 2021).

Analysis of the local channel density within synapses was performed by cluster analysis based on Voronoï tessellation constructed from Cav2.1 channel localizations analyzed with ThunderSTORM (ImageJ) and extracted local channel density using the software package SR-Tessler (Levet et al., 2015).

### Mathematical modelling

We performed computational model simulations to investigate the effect of Ca^2+^ channel locations and coupling distance on the release probability of a single synaptic vesicle (SV), shown in Figure 3. To do so, we performed deterministic simulations of calcium diffusion using the CalC (“Calcium Calculator”), version 6.8.6, by Victor Matveev (Allbritton et al., 1992; Chen et al., 2015; Delvendahl et al., 2015; Kobbersmed et al., 2020; Lou et al., 2005; Matveev et al., 2002; Schneggenburger and Neher, 2000; Tran and Stricker, 2018; Xu et al., 1997). All simulations were performed a box of 0.5 µm x 0.5 µm x 0.5µm with a grid size of 0.005 µm. All boundaries of the simulated volume were set to be reflective so that no Ca^2+^ would escape the volume. Parameter settings (see Table S6) for Ca^2+^ concentration and buffers were inspired by those used by Kobbersmed et al. (2020) but adapted to values used by previous CalC simulation studies on smaller bouton-sized synapses (Tran and Stricker, 2018). We did not model gating or Ca^2+^ influx of the channels and instead assumed Gaussian Ca^2+^ currents (Schneggenburger and Neher, 2000) with FWHM = 244 µs (Tran and Stricker, 2018) and peak at 2 ms after initiation of the simulation. The Ca^2+^ influx charge iCh was a free parameter (see Table S6) and adjusted such that the model reproduced the experimentally observed failure rates for typical numbers of calcium channels as observed here, assuming an open probability of 50%. In all simulations we registered the time-dependent Ca^2+^ concentration at coordinates (0.0, 0.0, 0.010 µm) as a proxy for the SV location. This Ca^2+^ concentration data was used as input for an allosteric model of Ca^2+^ activation of SV fusion by Lou et al. (2005). Parameters for the allosteric model were set as described by Lou et al. (2005) and the model was solved numerically. The release probability (pVr) of the SV in each simulation was then determined as the proportion of fused SVs (the fraction of 1) after 10 ms according to the allosteric model. To assess whether channels cooperate beyond simple statistical independence, we compared the simulated pVr with the pVr predicted assuming independent channel contributions:

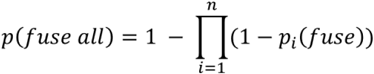

Locations of simulated Ca^2+^ channels were determined as follows: for Figure 3FCG, the closest channel was placed at a coupling distance of 30 nm from the SV. All other channels were placed with larger coupling distances which would by themselves cause very small pVrs (e.g. see the much lower pVr value for a channel at 100 nm vs. 30 nm in Fig. 3G). For Figure 3H-J, we placed N = 5, 10 or 15 Ca^2+^ channels at separate random locations (*dispersed* condition) or at the same random location (*clustered* condition) within the 0.5 µm x 0.5 µm bottom surface of the box. For Figure 3K-N, we simulated a *mixed* condition by placing a total of N = 10 channels with 5 of these placed randomly and independently (as in *dispersed*) and 5 in the same randomly picked location (as in *clustered*). We performed 90 simulations for each condition and value of N. For Figure S3, we placed the SV at (0,0), placed one channel 20 nm from the SV and varied the distance and angle of the second channel to the SV to investigate the effect of various combinations of channel positions.

To calculate failure rates for each condition (*mixed* and *clustered*) in Figure 3N, we compared each simulated release probability to a random number *r* ∼*U (0,1).* If pVr_i_ < r_i_, this was defined as release failure. This was done 10000 times for each condition to obtain a distribution of failure rates. Failure rate distributions were compared using a two-sample t-test, α = 0.05.

All analyses were done using Matlab (version R2024a).

### Statistical data analysis

Data analysis was mainly performed using GraphPad Prism 10 software. Data were compared using the parametric Student’s t-test, non-parametric Mann-Whitney test, Kolmogorov-Smirnov test, Chi-Squared test, and Correlation test. Data are represented as mean ± SEM or median with the interquartile range (IQR). Significance was evaluated at P<0.05, and the level of significance denoted by asterisks corresponds to: * p < 0.05; ** p < 0.01; *** p < 0.001; **** p < 0.0001.

The number of trajectories in SPT experiments, cell culture number (N), number of regions of interest (ROI, n) and number of cells analysed for individual experiments are described in the figure legends or related supplementary tables and figures. For comparisons between two groups, a parametric unpaired *t*-test was used, while multiple groups were analysed using one-way ANOVA followed by Tukey’s multiple comparisons test. For calculating the significance level GraphPad Prism 10 has been used. Error bars indicate the mean ± SEM if not stated different. The significance level is indicated as ****P < 0.0001, ***P < 0.001, *P < 0.1 within the figures and supplementary tables.

