## Supplementary Figures for "Functional relevance of mobile and clustered Ca_V_2.1 channels in central synapses"

**Supplementary Figure S1. Validation of constructs used within the study.**

**A)** Validation of Cre-recombinase mediated knock-down of $Ca_V2.1$ channels in hippocampal cultures probed with either anti-GFP nanobody or monoclonal anti-$Ca_V2.1$ antibody. Hippocampal neurons were infected with AAV-Cre virus at 3-4 DIV and labelled with the $Ca_V2.1$ targeting probes at 18-21 DIV, scale bar 20 µm. **B, C)** Normalized fluorescent intensities for anti-GFP nanobody and anti-$Ca_V2.1$ channels, data are from two independent cultures and 10 ROI for each condition. **D)** Schematic description of the mechanism for the controlled expression of intrabodies. **E)** Examples for the expression of transcription-controlled constructs in HEK-cells expressing either GFP on the outer leaflet of the membrane (GPI-GFP) or at the inner leaflet of the plasma membrane (myrGGFP), scale bar 5µm. **F, G)** Diffusion coefficient of Halo-tagged anti GFP intrabodies targeting Cacna1a$^{citrine}$ tagged channels expressing the transcription control unit (+CCR5) or not (ΔCCR5), represented as logarithmic frequency distribution or box blot. **H)** Distribution of the Mean jumping distance (MJD) for the synaptic population of $Ca_V2.1$ channels labelled with the transcription controlled intrabodies (gray) and not controlled intrabodies (black). **I)** Similar as in (H) for the axonal extrasynaptic population of $Ca_V2.1$ channels. Data in B-C are from 2 independent cultures and 10 ROIs for each, data in F-I are from 3 independent cultures. Data details and statistics are in supplementary Fig.1 table.

**Supplementary Figure S2. Fluorescent recovery after photo bleach for presynaptic $Ca_V2.1$ channels**

**A)** Example images of axons expressing the intrabody::Halo-tag to label presynaptic $Ca_V2.1$ channels. Clusters of $Ca_V2.1$ channels differ in their intensity and show a variability in the fluorescent recovery after photo bleach (FRAP) as indicated for three synapses (arrows) before bleaching (control), right after bleaching (bleach), and after steady state recovery (recovery). **B)** Time course of fluorescence recovery normalized to the initial fluorescence intensity. Data represents the average recovery rate from multiple boutons. **C)** Quantification of the recovery rate of endogenous $Ca_V2.1$ channels tagged with the intrabody. The mean recovery rate was about 0.32 ± 0.17, $n_{synapses}$ = 73, $n_{cultures}$ = 3). Data are presented as mean ± SEM. **D)** Example images for axons expressing cytosolic GCamp6f and Synaptophysin::mRFP to label presynaptic boutons and monitor potential calcium signalling along axons (scale bar 20 µm and 5µm for the enlarged region). Example calcium responses to a brief field stimulation (0.5 ms, 50 mA) within the synapse and 1 or 2 µm away from the centre. **F)** Average calcium responses within synapses, 1µm or 2µm away from synapses. Data details and statistics are in supplementary Fig.2 table

**Supplementary Figure S3**. **Effect on pVr for various combinations of locations of two $Ca^{2+}$ channels**

(**A**) Schematic representation of channel locations for each simulation. In all simulations, [$Ca^{2+}$] is registered at the simulated vesicle location (0,0) and the closest $Ca^{2+}$ channel (ch1) is placed at (0,0.020 µm). For each simulation, the second channel (ch2) is placed at various angles (black → light red) and at various distances (circle = 0.030 µm, diamond = 0.050 µm, square = 0.100 nm) around the SV location. (**B**) Effect of increased angle between ch2 and x-axis on the distance between ch1 and ch2, shown for each SV-ch2 distance ($d_2$). (**C**) Effect of increased angle (and thereby also inter-channel distance) for each $d_2$ on pVr.

**Supplementary Figure S4. Analysis of release properties and contribution of different calcium channel isoforms for glutamatergic synapses in primary hippocampal cultures**

**A)** Example image of transfected neurons expressing iGluSnFr3 and jRGECO1a::synaptophysin. **B)** Example traces for regions as indicated in A). **C, D, E)** Distribution of glutamate responses from axons and dendrites with and without TTX (1µM). **F, G)** Fraction of mono and multivesicular responses. **H)** Protocol of cumulative application of specific calcium channel blockers and corresponding synaptic calcium responses. **I)** Distribution of the fractions of $Ca_V2.1$ (P/Q), $Ca_V2.2$ (N) and $Ca_V1.2$ (L) channel sensitive synapses in primary hippocampal mouse neurons 18-23 DIV.

Data details and statistics are in supplementary Fig.4 table

**Supplementary Figure S5. Immunohistochemical labelling of pre- and postsynaptic markers plus $GABA_B$Rs in hippocampal synapses.**

Quantification of colocalization of Homer, Bassoon and GABABR. Example images for all three proteins are given, Bassoon (green), Homer (magenta), GABAB receptors (turquoise). Data are from three independent cultures out of 30 ROI. About 82% ± 10% of GABAB receptors colocalize in glutamatergic synapses. The scale bar represents 5µm.

**Supplementary Table 1 measurements**

| **Fig. 1** | Condition | $N_{roi/syn.}$ | $N_{cultures}$ | Mean | SEM |
|---|---|---|---|---|---|
| D | WT neurons + CCR5-IB | 6 | 2 | 1.000 | 0.163 |
| D | Cre neurons + CCR5-IB | 11 | 2 | 0.184 | 0.029 |
| E | WT neurons + ΔCCR5-IB | 10 | 2 | 1.000 | 0.137 |
| E | Cre neurons + ΔCCR5-IB | 8 | 2 | 0.854 | 0.156 |
| F | WT neurons + CRY2-olig-IB | 10 | 2 | 1.000 | 0.173 |
| F | Cre neurons + CRY2-olig-IB | 10 | 2 | 0.031 | 0.005 |
| G | Bassoon positive synapses | 23 | 3 | 0.608 | 0.019 |
| G | Inhibitory synapses | 20 | 3 | 0.857 | 0.026 |
| G | Excitatory synapses | 16 | 3 | 0.947 | 0.006 |
| H | Cav2.1 intensity in inhibitory synapses | 27 | 3 | 1.000 | 0.082 |
| H | Cav2.1 intensity in excitatory synapses | 30 | 3 | 1.085 | 0.140 |
| I | Occupancy rate – construct I | 11 | 3 | 0.476 | 0.040 |
| I | Occupancy rate – construct II | 30 | 3 | 0.756 | 0.016 |
| I | Occupancy rate – construct III | 29 | 3 | 0.819 | 0.019 |

**statistics**

| **Fig. 1** | Comparison | significance | p-value | Statistical method |
|---|---|---|---|---|
| D | WT + CCR5-IB vs. Cre + CCR5-IB | **** | <0.0001 | Unpaired t test |
| E | WT + ΔCCR5-IB vs. Cre + ΔCCR5-IB | ns | 0.412 | Unpaired t test |
| F | WT + CRY2-IB vs. Cre + CRY2-IB | **** | <0.0001 | Unpaired t test |
| H | Cav2.1 intensity in excitatory vs. inhibitory synapses | ns | 0.612 | Unpaired t test |

**Supplementary Table 2 measurements**

| **Fig. 2** | Condition | $n_{roi/syn.}$ | $N_{cultures}$ | $n_{trajectories}$ | Mean | SEM |
|---|---|---|---|---|---|---|
| F | Control | 189 | 3 | N/A | 2.241 | 0.128 |
| F | Intrabody | 135 | 3 | N/A | 2.303 | 0.077 |
| G | MSD of synaptic Cav2.1 | 205 | 4 | 4510 | N/A | N/A |
| G | MSD of extra-synaptic Cav2.1 | 184 | 4 | 5520 | N/A | N/A |
| H | Diff. Coeff. of synaptic Cav2.1 | 205 | 4 | 4510 | N/A | N/A |
| H | Diff. Coeff. of extra-synaptic Cav2.1 | 184 | 4 | 5520 | N/A | N/A |
| I | MJD of synaptic Cav2.1 | 300 | 5 | 105000 | N/A | N/A |
| I | MJD of extra-synaptic Cav2.1 | 260 | 5 | 91520 | N/A | N/A |
| J | MJD seg. of synaptic Cav2.1 | 300 | 5 | 105000 | N/A | N/A |

| | | | | | | |
|---|---|---|---|---|---|---|
| J | MJD seg. of extra-synaptic Cav2.1 | 260 | 5 | 91520 | N/A | N/A |
| K | Synaptic MJD seg. Of control | 98 | 3 | 47040 | N/A | N/A |
| K | Synaptic MJD seg. Of Lat A | 117 | 3 | 42000 | N/A | N/A |
| K | Synaptic MJD seg. Of control | 91 | 3 | 40950 | N/A | N/A |
| K | Synaptic MJD seg. Of Nocodazole | 126 | 3 | 47250 | N/A | N/A |
| L | Extra-synaptic MJD seg. of control | 91 | 3 | 40950 | N/A | N/A |
| L | Extra-synaptic MJD seg. of Lat A | 247 | 3 | 92625 | N/A | N/A |
| L | Extra-synaptic MJD seg. of control | 121 | 3 | 43560 | N/A | N/A |
| L | Extra-synaptic MJD seg. of Nocodazole | 136 | 3 | 70720 | N/A | N/A |
| M | Synaptic MJD seg. of control | 158 | 3 | 39500 | N/A | N/A |
| M | Synaptic MJD seg. of TTX | 168 | 3 | 42000 | N/A | N/A |
| M | Synaptic MJD seg. of Bicuculine | 167 | 3 | 52000 | N/A | N/A |
| M | Synaptic MJD seg. of BAPTA | 81 | 3 | 21060 | N/A | N/A |
| N | Extra-synaptic MJD seg. of control | 112 | 3 | 58240 | N/A | N/A |
| N | Extra-ynaptic MJD seg. of TTX | 135 | 3 | 75600 | N/A | N/A |
| N | Extra-synaptic MJD seg. of Bicuculine | 121 | 3 | 58080 | N/A | N/A |
| N | Extra-synaptic MJD seg. of BAPTA | 104 | 3 | 33280 | N/A | N/A |
| Q | Correlation failure rate number of channels | 78 | 3 | N/A | N/A | N/A |

**statistics**

| **Fig. 2** | Comparison | Significance | p-value | Statistical method |
|---|---|---|---|---|
| J | Syn. vs Extrasyn. | * | 0.0261 | Chi-square test |
| K | Syn. Control vs Syn. Lat A | ns | 0.653 | Chi-square test |
| K | Syn. Control vs Syn. Nocodaz. | ns | 0.386 | Chi-square test |
| L | extra-syn. Control vs Syn. Lat A | ** | 0.001 | Chi-square test |
| L | extra-syn. Control vs Syn. Nocodaz. | ** | 0.001 | Chi-square test |
| M | Syn. Control vs Syn. TTX | * | 0.0257 | Chi-square test |
| M | Syn. Control vs Syn. Bicuculine | ns | 0.757 | Chi-square test |
| M | Syn. Control vs Syn. BAPTA | ** | 0.006 | Chi-square test |
| N | extra-syn. Control vs Syn. TTX | ns | 0.911 | Chi-square test |
| N | extra-syn. Control vs Syn. Bicuculine | ns | 0.606 | Chi-square test |
| N | Extra-syn. Control vs Syn. BAPTA | ns | 0.379 | Chi-square test |
| Q | Correlation failure rate number of channels | **** | <0.0001 | Pearson |

**Supplementary Table 3 measurements/statistics**

| **Fig. 3** | Comparison | n | N | Median/IQR | Statistical test |
|---|---|---|---|---|---|
| D | Local surface/synapse | 1.683 | 2 | 0.041 $\mu m^2$; 0.022/0.083 | Mann-Whitney test |

| | | | | | |
|---|---|---|---|---|---|
| D | Local surface/nanocluster | 916 | 2 | 0.0054 $\mu m^2$; 0.0025/0.01 | **** |
| E | Local density/synapse | 1.683 | 2 | 0.024 loc./$nm^2$; 0.014/0.037 | Mann-Whitney test |
| E | Local density/nanocluster | 916 | 2 | 0.15 loc./$nm^2$ 0.08/0.19 | **** |

**Supplementary table 4 / measurements**

| Fig. 4 | Condition | $n_{roi/syn.}$ | $N_{cultures}$ | $n_{trajectories}$ | Mean | SEM |
|---|---|---|---|---|---|---|
| C | Synaptic $Ca^{2+}$ in control | 85 | 3 | N/A | 2.128 | 0.067 |
| C | Synaptic $Ca^{2+}$ in X-link | 85 | 3 | N/A | 2.109 | 0.030 |
| D | MJD seg. Of control | 72 | 3 | 19440 | N/A | N/A |
| D | MJD seg. Of X-link | 72 | 3 | 16560 | N/A | N/A |
| E | # of Cav2.1 channels in control | 79 | 3 | N/A | 29 | 3 |
| E | # of Cav2.1 channels in X-link | 86 | 3 | N/A | 30 | 3 |
| H | Cluster surface of control | 71 | 3 | N/A | 0.009 | 0.0006 |
| H | Cluster surface of X-link | 63 | 3 | N/A | 0.002 | 0.0002 |
| I | Cluster density of control | 71 | 3 | N/A | 0.02 | 0.001 |
| I | Cluster density of X-link | 63 | 3 | N/A | 0.27 | 0.020 |
| | Conditions | $n_{roi/syn.}$ | $N_{cultures}$ | $glutamate_{events}$ | median | IQR (25%/75%) |
| J | Frequency distribution Glutamate responses to 1 Hz stimuli intrabody | 25/964 | 7 | 30225 | 0.4 | 0.27/0.59 |
| J | Frequency distribution Glutamate responses to 1 Hz stimuli x-link | 20/891 | 7 | 27274 | 0.43 | 0.28/0.57 |
| K | Frequency distribution Glutamate responses to 1 Hz stimuli control | 29/541 | 9 | 23388 | 0.35 | 0.2/0.49 |
| K | Frequency distribution Glutamate responses to 1 Hz stimuli Cre-induced knock down of $Ca_V2.1$ | 16/215 | 5 | 9837 | 0.3 | 0.2/0.53 |
| L | Failure rate for control | 19/964 | 9 | N/A | 0.39 | 0.17/0.55 |
| L | Failure rate for Cre-induced knock-down | 21/891 | 5 | N/A | 0.49 | 0.41/0.65 |
| L | Failure rate for intrabody | 29/541 | 7 | N/A | 0.37 | 0.29/0.89 |
| L | Failure rate for x-link | 16/215 | 7 | N/A | 0.47 | 0.37/0.66 |
| N | Release area control | 200 | 4 | N/A | 0.032 | 0.028/0.038 |
| N | Release area Cre-induced knock-down | 211 | 4 | N/A | 0.03 | 0.027/0.032 |
| N | Release area intrabody | 64 | 4 | N/A | 0.028 | 0.025/0.032 |
| N | Release area x-link | 281 | 4 | N/A | 0.015 | 0.011/0.017 |
| | Conditions | $n_{roi/syn.}$ | $N_{cultures}$ | $burst_{events}$ | median | IQR (25%/75%) |
| P | 10 Hz intrabody | 12/119 | 3 | 119 | 113.4 | 75.7/199 |
| P | 10 Hz x-link | 21/230 | 3 | 230 | 140.6 | 91.5/221.7 |
| P | 20 Hz intrabody | 14/187 | 3 | 187 | 264.5 | 152/392 |
| P | 20 Hz x-link | 21/210 | 3 | 210 | 189.9 | 141/264 |
| P | 50 Hz intrabody | 11/137 | 3 | 137 | 297.4 | 204/436 |
| P | 50 Hz x-link | 16/152 | 3 | 152 | 226.8 | 144/284 |

**statistics**

| Fig. 4 | Comparison | Significance | p-value | Statistical method |
|---|---|---|---|---|
| C | Synaptic $Ca^{2+}$ in control vs. X-link | ns | 0.798 | Unpaired t test |
| D | MJD in control vs. X-link | ** | 0.0039 | Chi-square test |
| E | # of Cav2.1 in control vs. X-link | ns | 0.735 | Unpaired t test |
| H | Cluster surface in control vs. X-link | **** | <0.0001 | Unpaired t test |
| I | Cluster density in control vs. X-link | **** | <0.0001 | Unpaired t test |
| J | Frequency distribution glutamate amplitudes Intrabody versus x-link | ns | 0.7227 | Kolmogorov-Smirnov test |
| K | Frequency distribution glutamate amplitudes WT ($Cacna1a^{citrine}$) versus Cre-KD | ns | 0.2914 | Kolmogorov-Smirnov test |
| L | Failure rate | * | 0.0108 | One way ANOVA |
| L | Failure rate intrabody versus x-link | * | 0.0159 | Holm-Sidak multiple comparison test |
| L | Failure rate control versus Cre-KD | * | 0.0159 | Holm-Sidak multiple comparison test |
| N | Release area | **** | <0.0001 | Kruskal-Wallis test |
| N | Release area Control versus x-link | **** | <0.0001 | Dunn's multi comparison test |
| N | Release area Control versus intrabody | ns | 0.1678 | Dunn's multi comparison test |
| N | Release area Control versus Cre-KD | ns | 0.4001 | Dunn's multi comparison test |
| P | Area under the curve (AUC) | *** | <0.0001 | One way ANOVA |
| P | AUC 10Hz control (grey)/x-link (red) | ns | 0.2255 | Dunn's multi comparison test |
| P | AUC 20Hz control (grey)/x-link (red) | **** | <0.0001 | Dunn's multi comparison test |
| P | AUC 50Hz control (grey)/x-link (red) | **** | <0.0001 | Dunn's multi comparison test |

**Supplementary table 5 measurements**

| Fig. 5 | Condition | $n_{roi/syn.}$ | $N_{cultures}$ | $n_{trajectories}$ | Mean | SEM |
|---|---|---|---|---|---|---|
| B | Synaptic $Ca^{2+}$ in control | 775 | 3 | N/A | 3.496 | 0.255 |
| B | Synaptic $Ca^{2+}$ after baclofen | 612 | 3 | N/A | 2.347 | 0.118 |
| C | % of silenced syn. in WT | 17 | 3 | N/A | 63.66 | 4.99 |
| C | % of responsive syn. in WT | 17 | 3 | N/A | 36.34 | 4.99 |
| C | % of silenced syn. in WT + conotoxin | 12 | 3 | N/A | 66.16 | 5.26 |
| C | % of responsive syn. in WT+ conotoxin | 12 | 3 | N/A | 33.8 | 5.26 |

| | | | | | | |
|---|---|---|---|---|---|---|
| C | % of silenced syn. in Cav2.1 KO | 10 | 3 | N/A | 89.75 | 2.52 |
| C | % of responsive syn. in Cav2.1 KO | 10 | 3 | N/A | 10.25 | 2.52 |
| D, E | MJD of silenced synapses | 115 | 3 | 4179 | N/A | N/A |
| D, E | MJD of responsive synapses | 135 | 3 | 2643 | N/A | N/A |
| | condition | $n_{ROI/syn.}$ | $N_{cultures}$ | $glutamate_{events}$ | median | IQR (25%/75%) |
| G | Frequency distribution Glutamate responses to 1 Hz stimuli intrabody | 21/784 | 4 | 30.473 | 0.44 | 0.3/0.62 |
| G | Frequency distribution Glutamate responses to 1 Hz stimuli intrabody+Baclofen | 20/347 | 4 | 4.206 | 0.23 | 0.18/0.31 |
| H | Failure rate intrabody | | 4 | N/A | 0.38 | 0.29/0.56 |
| H | Failure rate baclofen | 19/324 | 4 | N/A | 0.85 | 0.8/0.89 |
| H | Failure rate x-link | | 4 | N/A | 0.47 | 0.38/0.62 |
| H | Failure rate x-link+baclofen | 22/598 | 4 | N/A | 0.71 | 0.58/0.86 |
| J | Frequency distribution Glutamate responses to 1 Hz stimuli x-link | 16/766 | 4 | 25.396 | 0.47 | 0.4/0.6 |
| J | Frequency distribution Glutamate responses to 1 Hz stimuli x-link+baclofen | 22/574 | 4 | 9.033 | 0.21 | 0.17/0.28 |
| K | Release area intrabody | 89 | 4 | N/A | 0.027 | 0.015/0.041 |
| K | Release area baclofen | 22 | 4 | N/A | 0.006 | 0.003/0.009 |
| K | Release area x-link | 141 | 4 | N/A | 0.012 | 0.007/0.024 |
| K | Release area x-link+baclofen | 47 | 4 | N/A | 0.014 | 0.008/0.021 |

**statistics**

| **Fig. 5** | Comparison | Significance | p-value | Statistical method |
|---|---|---|---|---|
| B | Synaptic Ca$^{2+}$ in control vs. baclofen | *** | 0.0009 | Unpaired t test |
| C | % of silent/responding synapses in WT vs Ca$_V$2.1 KO | **** | <0.0001 | Chi-Square test |
| E | Silenced vs responsive syn. (MJD) | **** | <0.0001 | Chi-square test |
| G | Frequency distribution Glutamate responses to 1 Hz intrabody (grey) versus intrabody+baclofen (green) | **** | <0.0001 | Kolmogorov-Smirnov test |
| G | Fraction of mono-vesicular release | *** | 0.0002 | Unpaired t-test with Welch correction |
| H | Failure rate | **** | <0.0001 | One way ANOVA |

| | | | | |
|---|---|---|---|---|
| H | Failure rate intrabody versus Cry2 | ns | 0.0959 | Holm-Sidak multiple comparison test |
| H | Failure rate baclofen versus x-link+baclofen | * | 0.048 | Holm-Sidak multiple comparison test |
| H | Failure rate x-link versus x-link+baclofen | *** | 0.0001 | Holm-Sidak multiple comparison test |
| H | Failure rate intrabody versus baclofen | **** | 0.0001 | Holm-Sidak multiple comparison test |
| J | Frequency distribution Glutamate responses to 1 Hz x-link (red) versus x-link+baclofen (red/green stripes) | **** | <0.0001 | Kolmogorov-Smirnov test |
| J | Fraction of mono-vesicular release | ** | 0.0016 | Unpaired t-test with Welch correction |
| K | Release area | **** | <0.0001 | Kruskal-Wallis test |
| K | Release area intrabody versus baclofen | **** | <0.0001 | Dunn's multi comparison test |
| K | Release area baclofen versus x-link | *** | 0.0001 | Dunn's multi comparison test |
| K | Release area baclofen versus x-link+baclofen | *** | 0.0007 | Dunn's multi comparison test |

**Supplementary Figure 1, measurements**

| **Suppl. Fig.1** | Condition | $n_{roi}$ | $N_{cultures}$ | Cluster number | Cluster intensity | Statistical test |
|---|---|---|---|---|---|---|
| C | $Ca_V$2.1 cluster control | 20 | 2 | 1.02 ± 0.03 | 1.01 ± 0.03 | Mann-Whitney test ****; p<0.0001 |
| C | $Ca_V$2.1 cluster Cre-induced knock-down | 20 | 2 | 0.135 ± 0.02 | 0.6 ± 0.035 | Mann-Whitney test ****; p<0.0001 |
| | Condition | $n_{roi/syn.}$ | $N_{cultures}$ | $n_{trajectories}$ | Median | IQR |
| F | Diff. Coeff. of +CCR5 | 308 | 3 | 1332 | N/A | N/A |
| F | Diff. Coeff. of ΔCCR5 | 204 | 2 | 2973 | N/A | N/A |
| G | Diff. Coeff. of +CCR5 | 25 | 3 | N/A | 0.028 | [0.021, 0.031] |
| G | Diff. Coeff. of ΔCCR5 | 17 | 2 | N/A | 0.039 | [0.032, 0.050] |
| H | Syn. MJD of +CCR5 | 132 | 3 | 34929 | N/A | N/A |
| H | Syn. MJD of ΔCCR5 | 108 | 2 | 2426 | N/A | N/A |
| I | Axonal MJD of +CCR5 | 130 | 3 | 12563 | N/A | N/A |
| I | Axonal MJD of ΔCCR5 | 117 | 2 | 2920 | N/A | N/A |

**Supplementary Figure 1, statistics**

| **Suppl. Fig.1** | Comparison | Significance | p-value | Statistical method |
|---|---|---|---|---|
| C | $Ca_V2.1$ control versus Cre-KD | **** | <0.0001 | Mann-Whitney test |
| G | Diff. Coeff. of +CCR5 vs. ΔCCR5 | ** | 0.009 | Mann-Whitney test |

**Supplementary Figure 2, measurements**

| Suppl. Fig.2 | Condition | $n_{roi/syn.}$ | $N_{cultures}$ | Mean | SEM |
|---|---|---|---|---|---|
| B | synaptic Recovery over time | 73 | 3 | N/A | N/A |
| C | Recovery rate | 73 | 3 | 0.322 | 0.020 |

**Supplementary Figure 4, measurements**

| **Suppl.Fig.4** | Condition | $N_{cultures}$ | $n_{events}$ | median | IQR |
|---|---|---|---|---|---|
| C | iGluSnFR3 axonal 1µM TTX | 4 | 1243 | 0.21 | 0.13/0.27 |
| D | iGluSnFR3 axonal | 5 | 2072 | 0.4 | 0.2/0.85 |
| E | iGluSnFR3 dendritic | 7 | 9507 | 0.97 | 0.42/1.7 |
| | Condition | $N_{cultures}$ | ROI | $mean_{normalized}$ | SEM |
| F | multi-vesicular release TTX | 4 | 14 | 0.31 | 0.045 |
| F | multi-vesicular release axon | 5 | 10 | 0.52 | 0.07 |
| F | multi-vesicular release dendrite | 7 | 7 | 0.71 | 0.08 |
| G | mono-vesicular release TTX | 4 | 14 | 0.69 | 0.045 |
| G | mono-vesicular release axon | 5 | 10 | 0.48 | 0.07 |
| G | mono-vesicular release dendrite | 7 | 7 | 0.28 | 0.08 |
